# Coping with oxidative stress in extreme environments: the distinctive roles played by *Acinetobacter* sp. Ver3 superoxide dismutases

**DOI:** 10.1101/2021.10.11.463953

**Authors:** Bruno Alejandro Steimbrüch, Mariana G. Sartorio, Néstor Cortez, Daniela Albanesi, María-Natalia Lisa, Guillermo Daniel Repizo

## Abstract

*Acinetobacter sp*. Ver3 is a polyextremophilic strain characterized by a high tolerance to radiation and pro-oxidants. The Ver3 genome comprises the *sodB* and *sodC* genes encoding an iron (^AV3^SodB) and a copper/zinc superoxide dismutase (^AV3^SodC), respectively; however, the specific role(s) of these genes has remained elusive. We show that the expression of *sodB* remained unaltered in different oxidative stress conditions whereas *sodC* was up-regulated in the presence of blue light. Besides, we studied the changes in the in vitro activity of each SOD enzyme in response to diverse agents and solved the crystal structure of ^AV3^SodB at 1.34 Å, one of the highest resolutions achieved for a SOD. Cell fractionation studies interestingly revealed that ^AV3^SodB is located in the cytosol whereas ^AV3^SodC is also found in the periplasm. Consistently, a bioinformatic analysis of the genomes of 53 *Acinetobacter* species pointed out the presence of at least one SOD type in each compartment, suggesting that these enzymes are separately required to cope with oxidative stress. Surprisingly, ^AV3^SodC was found in an active state also in outer membrane vesicles, probably exerting a protective role. Overall, our multidisciplinary approach highlights the relevance of SOD enzymes when *Acinetobacter* spp. are confronted with oxidizing agents.

## INTRODUCTION

High-altitude Andean lakes (HAALs) along the central Andes area in South America undergo extreme environmental conditions such as high concentrations of salts and metalloids, wide daily temperature variations and high exposure to UV radiation^1^. These ecosystems are thus sources of extremophile microorganisms that evolved diverse biological strategies to survive hostile environments. Indeed, about 1000 bacterial strains have been isolated from the area to constitute the Extremophile Culture Collection from HAALs^2,3^.

*Acinetobacter sp*. Ver3 and Ver7 are two phylogenetically related strains that were isolated from Laguna Verde located at 4,400 m above sea level. These strains are better adapted to survive exposure to UV-B radiation compared to the collection strains *A. baumannii, A. lwoffii* and *A. johnsonii*^4,5^ and also display high tolerance to the chemical pro-oxidants H_2_O_2_ and methyl viologen (MV)^5^. Available evidence indicates that photolyases might be involved in an efficient DNA repair system that contributes to the UV-B resistance phenotype of the Andean *Acinetobacter* isolates^4,6^. On the other hand, the high tolerance to pro-oxidants of these strains has led to the investigation of their catalases and superoxide dismutases (SODs), since they are the most important enzymes for the elimination of reactive oxygen species (ROS) resulting from partial reduction of oxygen in aerobic cells. The catalase KatE1 from *Acinetobacter sp*. Ver3 (^AV3^KatE1) has one of the highest catalytic activities reported for a catalase^7^. This enzyme is constitutively expressed in high amounts in the bacterial cytosol and acts as the main protecting catalase against H_2_O_2_ and UV-B radiation^5,8^. Instead, ^AV3^KatE2 is a periplasmic enzyme that is strongly induced by peroxide and UV radiation and provides additional protection against pro-oxidants^8^. H_2_O_2_ is formed as a by-product of the respiratory electron transport chain through the reduction of molecular dioxygen or by disproportionation of the superoxide ion catalysed by SODs. Relatively little is known about the specific functions of SOD variants in *Acinetobacter* species.

SODs are widely distributed diverse metalloenzymes that are classified according to the metal cofactor present in the active site. Manganese SODs (MnSODs or SodA enzymes) and iron SODs (FeSODs or SodB enzymes) are found in the bacterial cytosol and have very similar active sites, suggesting that these enzymes are closely related and evolved from a common ancestor, a relation that is supported by the identification of SOD variants termed cambialistic that use either iron or manganese (cambialistic Fe/MnSODs) depending on metal availability^9,10^. Early studies showed that the inactivation of *sodA* and *sodB* in *E. coli* leads to increased susceptibility to oxidative stress, higher mutation rates and growth defects on minimal media due to the inactivation by O_2_ of enzymes involved in amino acid biosynthesis^11^. On the other hand, copper-zinc SODs (CuZnSODs or SodC enzymes) belong to a different lineage and are found in the periplasmic space of gram negative bacteria^12^. The inability of O_2_^-^ to cross the cytoplasmic membrane and the subcellular location of CuZnSODs suggests that periplasmic SODs most likely protect bacteria from exogenous O_2_^-13^. The physiological relevance of these enzymes was emphasized in studies of the pathogenic bacterium *Salmonella typhimurium*, which produces three periplasmic CuZnSODs. SodCI, the most relevant isoform for the prevention of oxidative damage in this species, is found in strains associated with non-typhoid *Salmonella* bacteremia^14–16^ and is necessary for protection against the oxidative burst of phagocytes and virulence^17,18^.

Diverse *sod* genes have been reported to coexist in *Acinetobacter* genomes^19,20^. For instance, *A. baumannii* ATCC 17978 encodes a gene *sodB* and a gene *sodC*. A *sodB* mutation in this strain leads to increased susceptibility to oxidative stress and the antibiotics colistin and tetracycline. Moreover, bacterial motility is affected and virulence is attenuated^19^. The transcription of *sodC* in the same strain was up-regulated in the presence of copper and zinc ions. However, the total SOD activity in exponentially growing cultures remained unchanged even though these metal ions contribute to bacterial resistance to ROS^20^.

To better understand the contributions of different types of SOD enzymes to resistance to oxidative stress, we conducted a multidisciplinary study of SODs present in an extremophile *Acinetobacter* species. Thus, we have characterized the enzymes FeSOD and CuZnSOD from the polyextremophile *Acinetobacter sp*. Ver3 (^AV3^SodB and ^AV3^SodC, respectively) by biochemical and structural methods. Besides, we present evidence that ^AV3^SodB is a cytosolic enzyme whereas ^AV3^SodC is directed to the bacterial periplasm. Furthermore, we show that the transcription of *sodC* in this strain is triggered in response to blue light. Finally, we include a phylogenetic analysis of *Acinetobacter* SOD enzymes and provide grounds for further studies relevant to biotechnology and health fields.

## METHODS

### Bacterial strains, plasmids, and culture media

Bacterial strains and plasmids used in this work are listed in Table S1. All strains were grown in Luria Bertani (LB) broth, supplemented with 1.5 % w/v agar for solid medium when necessary. The antibiotics ampicillin (100 μg/ml), kanamycin (40 μg/ml), and chloramphenicol (20 μg/ml) were added for selection as required. *E. coli* cells were grown at 37 °C unless otherwise indicated. *Acinetobacter* strains were grown at 30 °C.

### DNA manipulation procedures

*Acinetobacter* sp. Ver3 genomic DNA was isolated following the CTAB method^21^. The ^AV3^SodB coding sequence was PCR-amplified using primers FMSOD3228F and FMSOD3228R (Table S2), digested with *Nco*I and *Sac*I and ligated into the corresponding sites in the previously designed pET3228 expression plasmid^22^, leading to vector pEV3SodB. The coding sequence for ^AV3^SodC devoid of its signal peptide (^AV3^SodC^-p^) was PCR-amplified using primers CSOD^-p^32F and CSOD^-p^32R, digested with *Bam*HI and *Xho*I and ligated into the corresponding sites of a modified version of the plasmid pET32a lacking the enterokinase cleavage site, leading to vector pEVSodC^-p^. This plasmid allows the production of the recombinant protein as a translational fusion to thioredoxin and a His_6_ tag.

All DNA digestions were performed following the enzyme manufacturer’s instructions. Constructions were verified by automated DNA sequencing.

### Protein production and purification

The plasmid pEVSodB was used to transform the SOD-deficient *E. coli* strain QC774(DE3) and produce recombinant ^AV3^SodB as follows. Transformed cells were grown at 37 °C to an OD_600nm_ of 0.6 in LB broth supplemented with ampicillin, kanamycin and chloramphenicol. Expression of ^AV3^SodB was induced by incubation with 0.05 mM IPTG during 6 h at 180 r.p.m. Cells were harvested (4,000 g at 4 °C, 15 min), resuspended in disruption buffer containing 50 mM Tris-HCl, 0.1 mM EDTA, 50 mM NaCl, pH 8.0, supplemented with 0.5 mM phenylmethylsulfonyl fluoride protease inhibitor (PMSF), 0.5 mM MgCl_2_ and 100 μg DNase per liter of culture, and lysed by sonication. The suspension was cleared by centrifugation at 4 °C and 17,000 g for 30 min and the supernatant was subjected to (NH_4_)_2_SO_4_ precipitation at 60 % w/v saturation. Precipitated proteins were collected by centrifugation (12,000 g at 4 °C, 15 min), the pellet was dissolved in 50 mM Tris-HCl, pH 8.0, and the suspension was dialyzed against 50 mM Tris-HCl, 50 mM NaCl, pH 8.0. The solution was loaded onto a Q Sepharose ion-exchange chromatography column (APbiotech) equilibrated with 50 mM Tris-HCl, pH 8.0. The enzyme was eluted by increasing the ionic strength of the buffer. Purified ^AV3^SodB eluted at 100 mM NaCl.

*E. coli* QC774(DE3) cells were employed for the production of recombinant ^AV3^SodC devoid of its signal peptide (^AV3^SodC^-p^) using the plasmid pEVSodC^-p^. Transformed cells were cultured at 37 °C to an OD_600nm_ of 0.6 in LB broth supplemented with ampicillin, kanamycin and chloramphenicol. Protein expression was induced by incubation with 0.05 mM IPTG during 6 h at 180 r.p.m. Cells were harvested (4,000 g at 4 °C, 15 min), resuspended in disruption buffer containing 50 mM Tris-HCl, 0.1 mM EDTA, 50 mM NaCl, pH 8.0, supplemented with 0.5 mM PMSF, 0.5 mM MgCl_2_ and 100 μg DNase per liter of culture, and lysed by sonication. The suspension was cleared by centrifugation at 4 °C and 17,000 g for 30 min and the supernatant was loaded onto a Ni-NTA column (Invitrogen) equilibrated with buffer 50 mM Tris-HCl, 50 mM NaCl, 5 mM imidazole, pH 8.0. After thoroughly washing the loaded column, the recombinant protein was eluted using the same buffer containing 100 mM imidazole. The thioredoxin and His_6_ tags were then removed by cleavage with thrombin during 3 h at 30 °C. The reaction was stopped by adding 4.4 mM EDTA and pure ^AV3^SodC^-p^ was finally obtained after a second step of affinity chromatography using a Ni-NTA column (Invitrogen).

### Protein analyses

Protein purification steps and subcellular fractionations were followed by SDS-PAGE by using the method of Laemmli^23^ for 12 % w/v acrylamide gels and Coomassie blue staining. Protein concentration was determined by the Bradford method^24^ using bovine serum albumin as standard.

Antibodies for ^AV3^SodB and ^AV3^SodC^-p^ were obtained by two consecutive injections of rabbits with 0.3 mg of purified proteins. In each case, the first subcutaneous injection was carried out with a 1:1 emulsion of the protein with Freund’s complete adjuvant. In the second inoculation, Freund’s incomplete adjuvant was used instead.

For immunoblot analysis, proteins were transferred to nitrocellulose membranes. Alkaline phosphatase-conjugated goat anti-rabbit IgG was employed as a secondary antibody (Sigma-Aldrich^R^, St. Louis, MI, USA). The antigen–antibody complex was detected by the alkaline phosphatase reaction, employing 5-bromo-4-chloro-3-indolyl-phosphate (BCIP) and nitro blue tetrazolium (NBT) as substrates (Roche^R^, Roche Applied Sciences, Indianapolis, IN, USA).

### SOD activity measurements

SOD activity was visualized *in situ* after electrophoresis of cellular lysates or the purified enzymes in nondenaturing polyacrylamide gels, as previously described^25^.

SOD activity was also determined spectrophotometrically by inhibition of the xanthine/xanthine oxidase-induced reduction of cytochrome *c*^26^, using a Cary WinUV UV-visible spectrophotometer equipped with a Cary Dual Cell Peltier accessory (Agilent Technologies, Santa Clara, CA, USA). Reactions were carried out at 25 °C, in 50 mM sodium phosphate buffer pH 7.8, using 210 pg/ml ^AV3^SodB or 930 pg/ml ^AV3^SodC^-p^.

### Thermal stability, pH tolerance and response to chemical agents of ^AV3^SodB and ^AV3^SodC

Treatments were applied on 0.5 mg/ml enzyme samples and residual activities were then measured spectrophotometrically, as described above. In each case, the activity of the untreated enzyme was defined as 100 %.

The thermostability of ^AV3^SodB and ^AV3^SodC^-p^ was assessed after heat treatments in 50 mM sodium phosphate buffer pH 7.8. ^AV3^SodB was incubated for 15, 30 or 45 min at 50, 55 or 60 °C. ^AV3^SodC^-p^ was incubated, instead, for 15, 30 or 45 min at 40, 45 or 50 °C.

The pH tolerance of ^AV3^SodB and ^AV3^SodC^-p^ was determined by incubating the enzymes in buffers with different pH values at 25 °C for 1 h. The buffer systems employed were 50 mM citrate (pH 3.0-6.0), Tris-HCl (pH 7.0-8.0), and glycine-NaOH (pH 9.0-12.0).

The effects of ethylenediaminetetraacetic acid (EDTA) and β-mercaptoethanol (BME) on SOD activity were determined at final concentrations of 1 mM or 10 mM of the compounds. The effects of detergents were investigated by using sodium dodecyl sulphate (SDS) and Tween 20 at final concentrations of 0.1 % v/v or 1 % v/v. The effects of denaturants were examined by using urea and guanidine hydrochloride at final concentrations of 8 M and 2.5 M, respectively. To test the stability in an organic medium, enzymes were incubated in the presence of ethanol at final concentrations of 20 % v/v or 50 % v/v. In each case, the enzyme was incubated with the chemical agent at 25 °C for 1 h in 50 mM sodium phosphate buffer pH 7.8.

### Crystallization, data collection and structure determination

Crystallization screenings were carried out using the sitting-drop vapor diffusion method and a Gryphon (Art Robbins Instruments) nanoliter-dispensing crystallization robot. Following optimization, crystals of ^AV3^SodB grew after 10-15 days from a 10 mg/ml protein solution, by mixing 1 μl of protein solution and 1 μl of mother liquor, in a hanging drop setup with 1 ml mother liquor in the reservoir, at 20 °C. Isomorphous diffraction-quality ^AV3^SodB crystals grew in 22 % w/v PEG 8000, 100 mM sodium cacodylate pH 7.5, 200 mM magnesium acetate, 1 mM flavin mononucleotide (FMN) or 30 % w/v PEG 4000, 100 mM sodium acetate pH 4.6, 200 mM ammonium acetate, 1 mM FMN or 30 % w/v PEG 4000, 100 mM Tris pH 8.5, 200 sodium acetate, 1 mM FMN as mother liquor. Single crystals were cryoprotected in mother liquor containing 25 % v/v glycerol and flash-frozen in liquid nitrogen. X-ray diffraction data were collected at the synchrotron beamline I04 (Diamond Synchrotron, UK), at 100 K, using wavelength 0.9795 Å. Diffraction data were processed using XDS^27^ and scaled with Aimless^28^ from the CCP4 program suite^29^.

The crystal structure of ^AV3^SodB was solved by molecular replacement using the program Phaser^30^ and the atomic coordinates of *E. coli* FeSOD from PDB entry 1ISA as search probe. The structure was refined through iterative cycles of manual model building with *Coot*^31^ and reciprocal space refinement with phenix.refine^32^. The iron atom and the FMN molecule were manually placed in a *mFo–DFc* sigma-A-weighted electron density map employing *Coot*^31^ and the resulting model was refined as described above. The final structure was validated through the Molprobity server^33^. It contained more than 97 % of residues within favoured regions of the Ramachandran plot, with no outliers. Figures were generated and rendered with Pymol 1.8.x (Schrödinger, LLC). Atomic coordinates and structure factors have been deposited in the Protein Data Bank under the accession code 7SBH.

### Subcellular fractionation

The cellular fractionation of *Acinetobacter* sp. Ver3 was carried out by a two-step osmotic shock process. Briefly, cells were harvested and resuspended in buffer 20 mM Tris-HCl, 20 % w/v sucrose, pH 8.0, supplemented with 0.5 mM PMSF. The resuspension volume was normalized according to the formula *V* = 0.05 OD_600_ *V*_c_, where *V*_c_ is the starting volume of the culture. The suspension was incubated 1 h at 4 °C. Then, 5 ml of ice-cold water were added per ml of suspension and the mixture was further incubated for 1 h under the same conditions. After centrifugation at 15,000 rpm for 30 min at 4 °C, the supernatant (*i.e*. the total periplasmic fraction) was collected and stored at 4 °C. The pellet, consisting of spheroplasts, was resuspended in an equivalent volume of disruption buffer and lysed by sonication as described above. Finally the total periplasmic fraction was centrifuged at 45,000 rpm for 3 h at 4°C allowing to obtain the soluble and insoluble periplasmic fractions. The latter was resuspended in 20 mM Tris-HCl pH 8.0.

### Purification of outer membrane vesicles (OMVs)

OMVs were purified from long-term stationary phase cultures of *Acinetobacter* sp. Ver3. Briefly, 1 ml of saturated culture was inoculated on 100 ml LB media and incubated overnight at 30 °C. Cells were then harvested and the supernatant was filtered through a 0.22 μm-membrane (Millipore) and centrifuged at 60,000 rpm for 4 h at 4 °C. The pellet containing the OMVs was resuspended in 20 mM Tris-HCl pH 8.0 and stored at −20 °C.

### RNA extraction and quantitative real-time reverse transcription PCR

Total RNA from *Acinetobacter sp*. Ver3 was isolated using TRI-Reagent^R^ (Molecular Research Center, Inc., Cincinnati, OH, USA) according to the manufacturer’s instructions. The quality and quantity of RNA samples were evaluated by agarose gel electrophoresis and electronic absorption (Abs_260nm/280nm_). Samples were treated with RQ1 RNase-free DNase (Promega, Madison, WI, USA) to remove possible DNA contaminants prior to reverse transcription. To obtain cDNA, 2 μg of RNA was used in the reverse transcription reaction with random primers, employing M-MLV Reverse Transcriptase (Promega^R^) according to the manufacturer’s instructions. Real-time PCRs were carried out on a StepOne device (Applied Biosystems, Thermo Fisher Scientific, Foster City, CA, USA) with 59 HOT FIREPol^R^ Eva-Green qPCR Mix Plus (ROX; Solis BioDyne, Tartu, Estonia) using specific primers (Table S2).

Results for ^AV3^*sodB* and ^AV3^*sodC* mRNAs were normalized to the mRNA of *recA* and *rpoB*, considered housekeeping genes, based on the standard curve quantitative method^34^. The specificity of each reaction was verified by melting curves between 55 °C and 95 °C with continuous fluorescence measurements.

### Bioinformatic analyses

The complete genome of *Acinetobacter sp*. Ver3 was previously reported and deposited in the RAST annotation server (https://rast.nmpdr.org/) and the NCBI database under accession number JFYL01^35^. The deduced protein sequences of *Acinetobacter sp*. Ver3 ^AV3^SodB and ^AV3^SodC (EZQ10255.1 and EZQ12222.1, respectively) were retrieved from the NCBI database. In each case, a multiple sequence alignment with related proteins was performed using MAFFT version 7.475 (https://mafft.cbrc.jp/alignment/software/) with default parameters.

Structurally characterized FeSODs, MnSODs and cambialistic Fe/MnSODs (here collectively referred to as Fe-MnSODs) available at the PDB with more than 30 % sequence identity compared to ^AV3^SodB and 80 % query coverage were sequence aligned with ^AV3^SodB. In the case of ^AV3^SodC, in addition to enzymes available at the PDB, 16 orthologous sequences retrieved from the NCBI with at least 57 % sequence identity with ^AV3^SodC and 70 % query coverage were also used. Alignments were visualized and annotated using Jalview^36^.

Two unrooted phylogenetic trees were built based on the amino acid sequence alignments of the *Acinetobacter sp*. Ver3 SOD enzymes by using the Maximum likelihood method (with a WAG+I+G4 substitution model). The bioinformatic software IQ-TREE multicore version 1.6.11^37^ was employed to generate both trees and their reliability was tested by bootstrapping with 10,000 repetitions. Results were visualized using the online toll iTOL V5.7 (https://itol.embl.de/).

A comparative genomic analysis of *Acinetobacter* strains available at the NCBI database on September 16^th^, 2020 was performed. Only strains with genomic sequence assemblies classified as complete or scaffolds were included in the study. The genomic and proteomic data corresponding to 314 strains were extracted, and a local database was constructed. SODs encoded by *A. baumannii* ATCC 17978 were used as query to perform BLASTp-sequence similarity searches^38^ against the local database, using 40 % sequence identity and 90 % query coverage cut-off values in the case of Fe-MnSODs and 55 % sequence identity and 75 % query coverage for CuZnSODs. The bioinformatic software Signal P v5.0 (http://www.cbs.dtu.dk/services/SignalP/) was used to predict the presence or absence of signal peptide sequences in Fe-MnSODs and CuZnSODs.

## RESULTS

### *Acinetobacter sp*. Ver3 encodes two putative SOD enzymes

The genome of *Acinetobacter sp*. Ver3 contains two putative SOD genes^35^. CL42_08295 is located on contig JFYL01000023.1, is the second gene within a putative two-gene operon and exhibits homology to *sodB* genes of gram-negative bacteria (Fig. 1). On the other hand, CL42_01680 is located on contig JFYL01000003.1, has homology to *sodC* genes of gram-negative bacteria and is predicted to constitute a monocistronic transcription unit.

**Figure 1.**
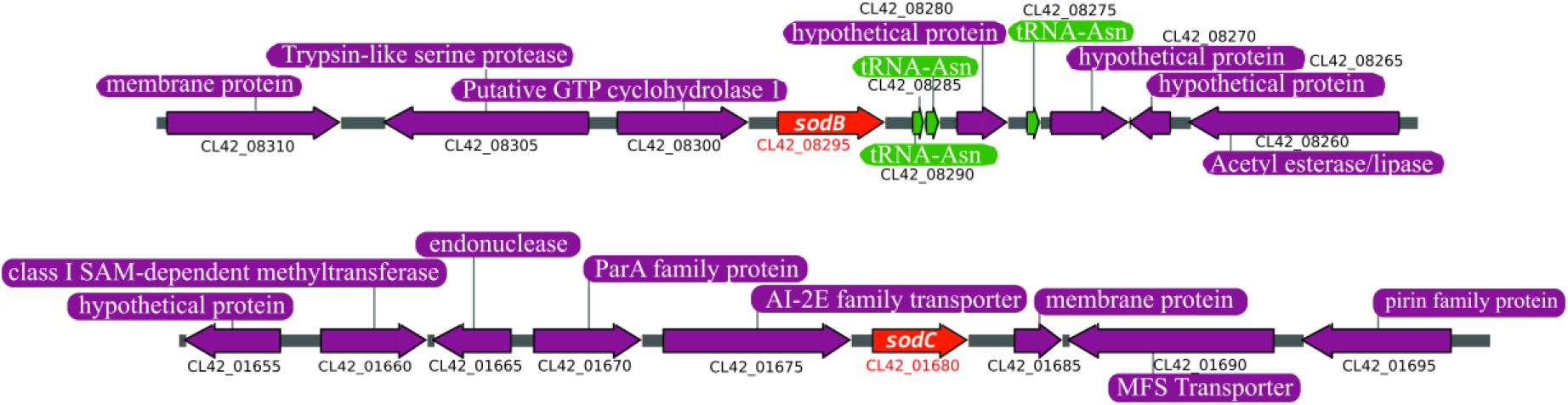
Environment of *sod* genes in *Acinetobacter sp*. Ver3. Schematic representation of the *Acinetobacter sp*. Ver3 *sodB* (up) and *sodC* (down) loci, present in contigs JFYL01000023 and JFYL01000003, respectively. *sod* genes are shown in red while proximal genes are displayed in purple. The locus tag is provided below each gene. Three tRNAs (green) are encoded in the *sodB* region.

The putative proteins encoded by the *sodB* and *sodC* genes from *Acinetobacter sp*. Ver3 (^AV3^SodB and ^AV3^SodC, respectively) were used as queries in BLASTp searches. The closest match of ^AV3^SodB in the PDB was the FeSOD from *Pseudomonas putida* (63 % identity, PDB code 3SDP). Fig. 2A displays a multiple sequence alignment of ^AV3^SodB and three structurally characterized related enzymes: the FeSOD (PDB code 1ISA) and MnSOD (PDB code 1VEW) from *E. coli* (60 % and 44 % amino acid identity, respectively) and the cambialistic SOD (PDB code 1QNN) from *P. gingivalis* (50 % identity). It shows that ^AV3^SodB harbours motifs that are characteristic of FeSODs, such as the motifs AAQ and DVEWHAYY involved in catalysis^39^. A phylogenetic tree built for ^AV3^SodB and diverse FeSODs, MnSODs as well as cambialistic Fe/MnSODs (here collectively referred to as Fe-MnSODs) of known structure consisted of three different clades, with archaeal and bacterial FeSODs grouped in two different clades whereas bacterial MnSODs constituted a third one (Fig. 2B). As expected, ^AV3^SodB clustered with bacterial FeSODs. On the other hand, the closest match of ^AV3^SodC in the PDB was the CuZnSOD from *Haemphylus ducrey* (53 % identity, PDB code 1Z9N). The sequence alignment in Fig. 2C highlights the conservation of metal binding residues in ^AV3^SodC and representative CuZnSODs retrieved from the NCBI database.

**Figure 2.**
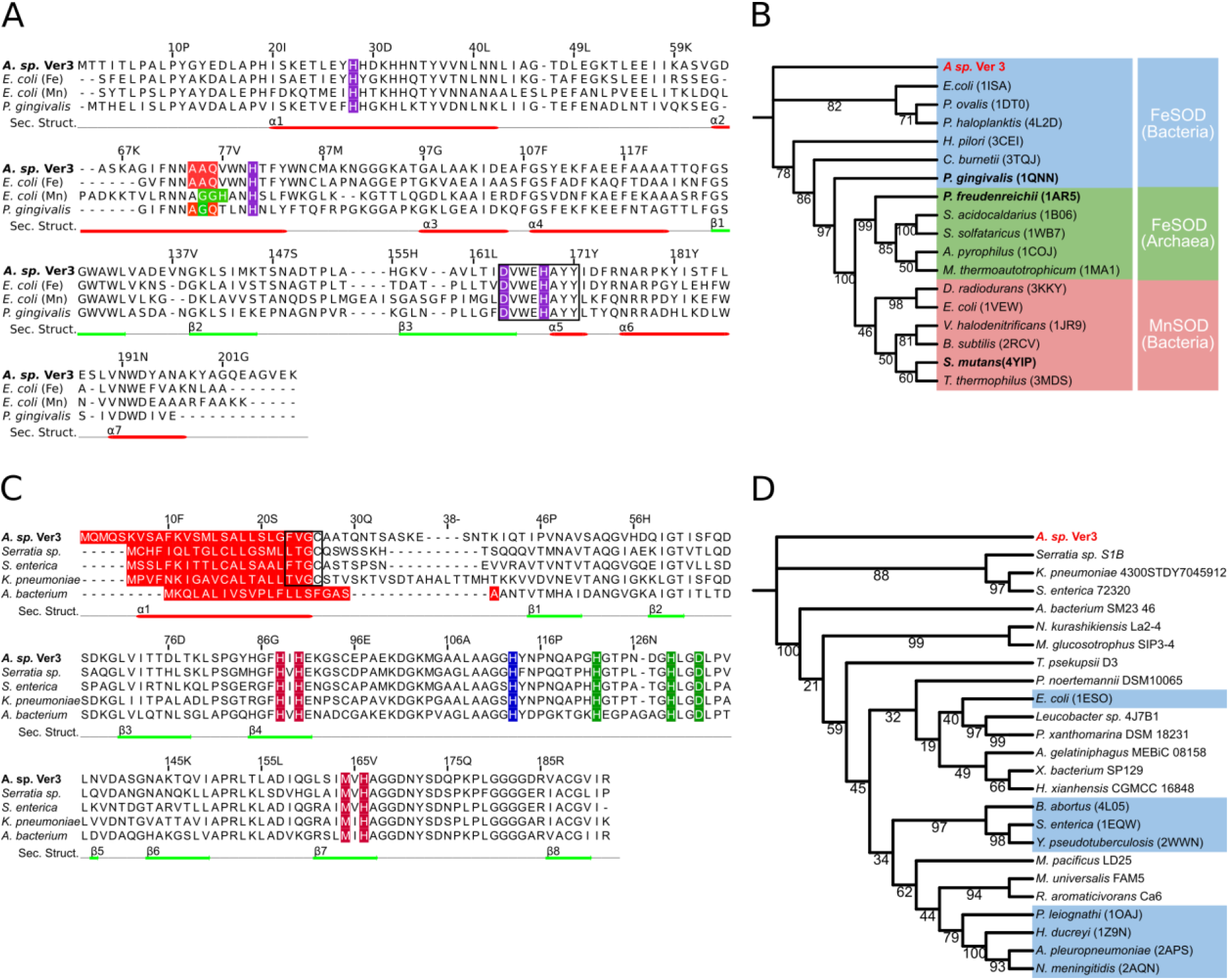
Sequence alignment and phylogenetic analysis of SOD enzymes. (A) Sequence alignment of FeSOD enzymes from selected species. Amino acids highlighted in purple correspond to conserved metal ligands. The motifs AAQ and GGH (in MnSODs, the *E. coli* enzyme is shown for comparison) are shown in red and green, respectively; the motif DVWEHAYYID comprising metal binding residues is boxed. Secondary structure elements observed in the crystal structure of ^AV3^SodB are depicted as red and green lines below the sequence alignment. (B) Phylogenetic tree of Fe-MnSODs of known structure. A clade of MnSODs (red) and two of FeSODs (blue and green) are distinguished. Cambialistics Fe/MnSODs are in bold. (C) Sequence alignment of CuZnSOD enzymes from selected species. Conserved metal ligands are shown in colour: copper ligands in vermillion, zinc ligands in green and a histidine residue that coordinates both metal ions in blue. Predicted signal peptides are signalled in red and lipobox sequences are boxed. Secondary structure elements predicted by Jalview^36^ are depicted as red and green lines below the sequence alignment. (D) Phylogenetic tree built for CuZnSOD sequences retrieved from the NCBI or the PDB (light blue boxes). Note that the signal peptide was removed from each sequence prior to alignment. Trees were constructed using the Maximum Likelihood algorithm; the robustness of the major branching points is indicated by the bootstrap values (10,000 repetitions).

Since FeSODs and CuZnSODs can present different subcellular locations^10^, we decided to analyse whether ^AV3^SodB and/or ^AV3^SodC presented signal sequences. No signal peptide was found for ^AV3^SodB using the Signal P 5.0 algorithm^40^, suggesting that this enzyme is likely cytosolic. In contrast, ^AV3^SodC displayed a 26-amino acid hydrophobic N-terminal sequence that is typical of signal peptides for protein export^41^ (Fig. 2C). A phylogenetic tree calculated from the predicted mature forms of ^AV3^SodC and a variety of CuZnSODs available at the PDB and the NCBI databases pointed out that ^AV3^SodC clusters with CuZnSODs from the gammaproteobacteria *Serratia sp., Salmonella enterica* and *Klebsiella pneumoniae* (Fig. 2D). Notably, enzymes belonging to this clade consistently display a putative lipoprotein signal peptide with a predicted lipobox cysteine residue^41,42^ (Fig. 2C).

### ^AV3^SodB and ^AV3^SodC^-p^ are active enzymes

To advance on their characterization, we decided to produce ^AV3^SodB and ^AV3^SodC by recombinant means. ^AV3^SodB was produced in a SOD-deficient *E. coli* strain and purified from the soluble cell fraction. The activity of ^AV3^SodB was assessed by *in situ* staining after nondenaturing PAGE, as previously described^25^. The obtained results (Fig. 3A) confirmed that ^AV3^SodB is an active enzyme. Besides, the electrophoretic mobility of the active ^AV3^SodB species matches that of the single SOD activity detected in soluble extracts of *Acinetobacter sp*. Ver3 (not shown).

**Figure 3.**
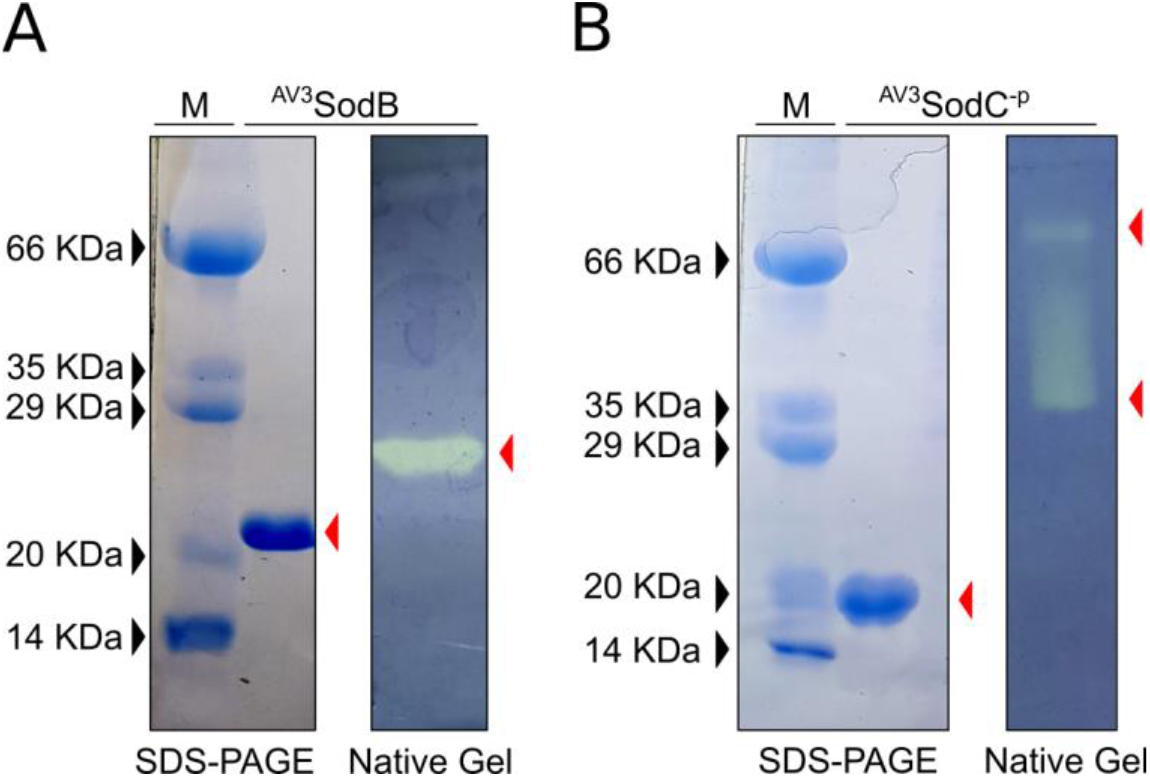
Recombinant Acinetobacter *sp*. Ver3 SODs. As isolated ^AV3^SodB (A) and ^AV3^SodC^-P^ (B) were analysed by SDS-PAGE and Coomassie Blue staining (left panels). SOD activity was revealed *in situ* in nondenaturing polyacrylamide gels (right panels). M: molecular weight marker.

We attempted to produce full-length ^AV3^SodC similarly to ^AV3^SodB, however these efforts repeatedly failed since multiple proteolytic fragments of the protein were found in SDS-PAGE analyses (not shown), suggesting a possible processing of the N-terminal peptide. Therefore, we assayed the production of a ^AV3^SodC truncated version lacking the 26-amino acid hydrophobic N-terminal sequence (^AV3^SodC^-p^). ^AV3^SodC^-p^ was successfully produced in *E. coli* and purified from the soluble cell fraction. ^AV3^SodC^-p^ exhibited two active species in non-denaturing gels, most likely corresponding to two different oligomeric states of the enzyme^10^ (Fig. 3B).

The specific activities of both *Acinetobacter sp*. Ver3 SOD enzymes were determined by the xanthine oxidase method (as described in the Methods section), resulting in values of 6,600 ± 200 U/mg and 1,800 ± 200 U/mg for ^AV3^SodB and ^AV3^SodC^-p^, respectively.

### Biochemical characterization of ^AV3^SodB and ^AV3^SodC^-p^

To evaluate the potential applications of *Acinetobacter* sp. Ver3 SODs in industry, we examined the effects of diverse physical and chemical effectors on enzyme activity. The thermostability of recombinant ^AV3^SodB and ^AV3^SodC^-p^ was investigated by pre-incubating the enzymes at different temperatures followed by measurements of the residual activity (Fig. 4A and B). Notably, the activity of ^AV3^SodB was virtually unaffected after a 45 min heat treatment at 50 °C. The enzyme even retained about 70 % activity after 45 min at 55 °C and 65 % activity after 30 min at 60 °C. On the other hand, ^AV3^SodC^-p^ conserved only 80 % and 40 % of its activity after incubation at 45 °C for 15 min or 45 min, respectively. Treatments at higher temperature resulted in a more pronounced loss of activity. These results indicated that ^AV3^SodB is more thermostable than ^AV3^SodC^-p^.

**Figure 4.**
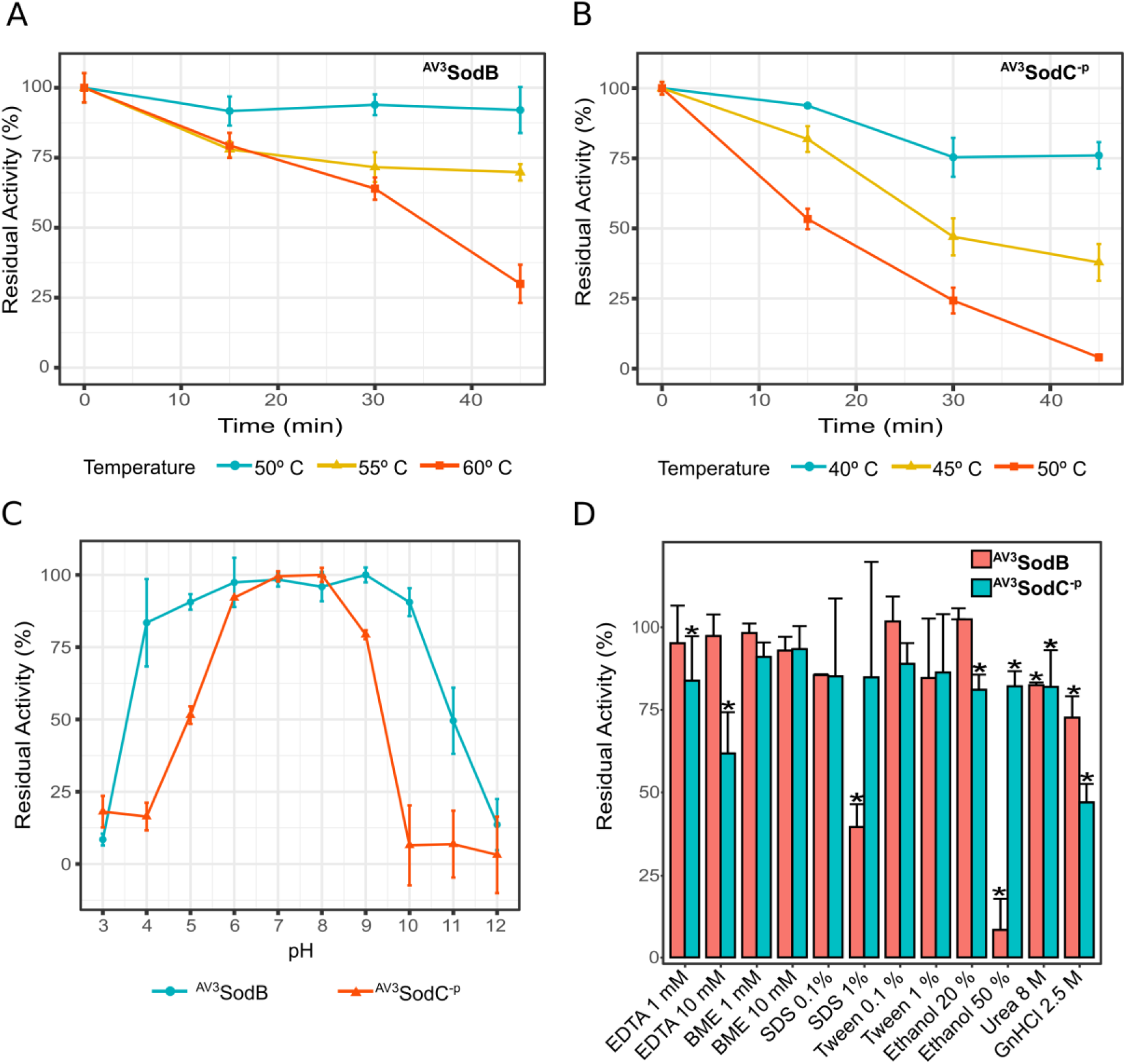
Biochemical characterization of *Acinetobacter sp*. Ver3 SODs. Thermostability of ^AV3^SodB (A) and ^AV3^SodC^-P^ (B). In each case, the residual SOD activity was measured after different heat treatments. (C) pH tolerance of ^AV3^SodB (blue line) and ^AV3^SodC^-P^ (red line). In each case, the residual SOD activity was measured at pH 7.8 after incubating the enzyme 1 h at 25° C in buffers with different pH values. (D) Effect of diverse chemical agents on ^AV3^SOD and ^AV3^SodC^-P^ activity. In each case, the residual SOD activity was measured after incubating the enzyme 1 h at 25° C with the compound. In all experiments, SOD activity was determined spectrophotometrically by inhibition of the xanthine/xanthine oxidase-induced reduction of cytochrome *c*, at 25° C; the activity of the untreated enzyme was defined as 100%. The reported values correspond to the mean of four measurements in three replicates of the experiment; bars indicate standard deviations.

The residual activity of ^AV3^SodB and ^AV3^SodC^-p^ was also measured after incubation in buffers with pH ranging from 3.0 to 12.0. ^AV3^SodB displayed a remarkable stability in the pH range 4.0-10.0, where it retained more than 75 % of its activity (Fig. 4C). ^AV3^SodC^-p^, on the other hand, conserved more than 75 % of activity in a narrower range, between pH values 6.0 and 9.0, pointing out that it is more sensitive to pH changes than ^AV3^SodB.

EDTA and β-mercaptoethanol (BME) (1 or 10 mM) were assayed as inhibitors of ^AV3^SodB and ^AV3^SodC^-p^ (Fig. 4D). ^AV3^SodB conserved more than 90 % of its activity in the presence of both compounds. However, while ^AV3^SodC^-p^ displayed high tolerance to BME, EDTA had an inhibitory effect, leading to a 40 % reduction of enzyme activity when used at 10 mM. SDS and Tween 20 (0.1% v/v or 1 % v/v) were used to investigate the influence of detergents on enzyme activity (Fig. 4D). SDS severely impaired ^AV3^SodB activity when used at 1 % v/v. All other effects were negligible. The denaturants urea and guanidine hydrochloride were also assessed, at final concentrations of 8 M and 2.5 M, respectively. ^AV3^SodC^-p^ lost approximately half of its activity when treated with guanidine hydrochloride. In all other cases enzyme activities remained higher than 70 %. To test the stability of *Acinetobacter* sp. Ver3 SODs in an organic solvent, the enzymes were incubated in reaction buffer supplemented with 20 % v/v or 50 % v/v ethanol. ^AV3^SodC^-p^ retained almost 80 % of its activity in both conditions while ^AV3^SodB lost nearly all its activity in 50 % v/v ethanol.

Overall, these results indicate a higher thermal stability and a broader pH range for ^AV3^SodB. Conversely, treatment with inhibitors showed mixed effects on enzyme activity, with ^AV3^SodB being altered mainly by SDS and ethanol, whereas ^AV3^SodC^-p^ was affected by EDTA and guanidine hydrochloride.

### Structural characterization of ^AV3^SodB

In order to explore the structural properties of ^AV3^SodB, we solved the crystal structure of the enzyme to 1.34 Å resolution (Table 1). ^AV3^SodB crystallized in space group C2221, with one protein molecule per asymmetric unit. The final atomic model comprises ^AV3^SodB residues 1-208. Two neighbouring ^AV3^SodB monomers related by crystallographic operations form a protein dimer with cyclic symmetry C2, resembling the quaternary structure of *E. coli* FeSOD (PDB code 1ISA) (Fig. 5A). The RMSD between the ^AV3^SodB monomer and the chain A in the *E. coli* FeSOD structure is 0.9 Å for 191 aligned residues. The ^AV3^SodB monomer adopts the two-domain fold conserved among FeSODs and MnSODs^10^, with a helical N-terminal domain and a C-terminal domain composed of three β-sheets surrounded by α-helices (Fig. 2A and Fig. 5A). Consistent with a dimeric functional state, the loop L2 (residues 43-68) that connects the two main helices in the N-terminal domain of ^AV3^SodB, the most variable in the primary sequence across species, unlike tetrameric enzymes^43,44^ is not extended and establishes hydrophobic interactions with the N-terminal loop L1 (residues 1-19) and the C-terminal domain (Fig. 2A and Fig. 5A). The interfacial area between ^AV3^SodB monomers is *ca*. 920 A^2^ and the residues involved in hydrogen bonds or salt bridges between subunits are Glu23, His32, Phe125, Ser127, Asn148, Glu167, His168, Tyr171 and Arg175, all conserved in *E. coli* FeSOD (Fig. 2A and Fig. 5A).

**Figure 5.**
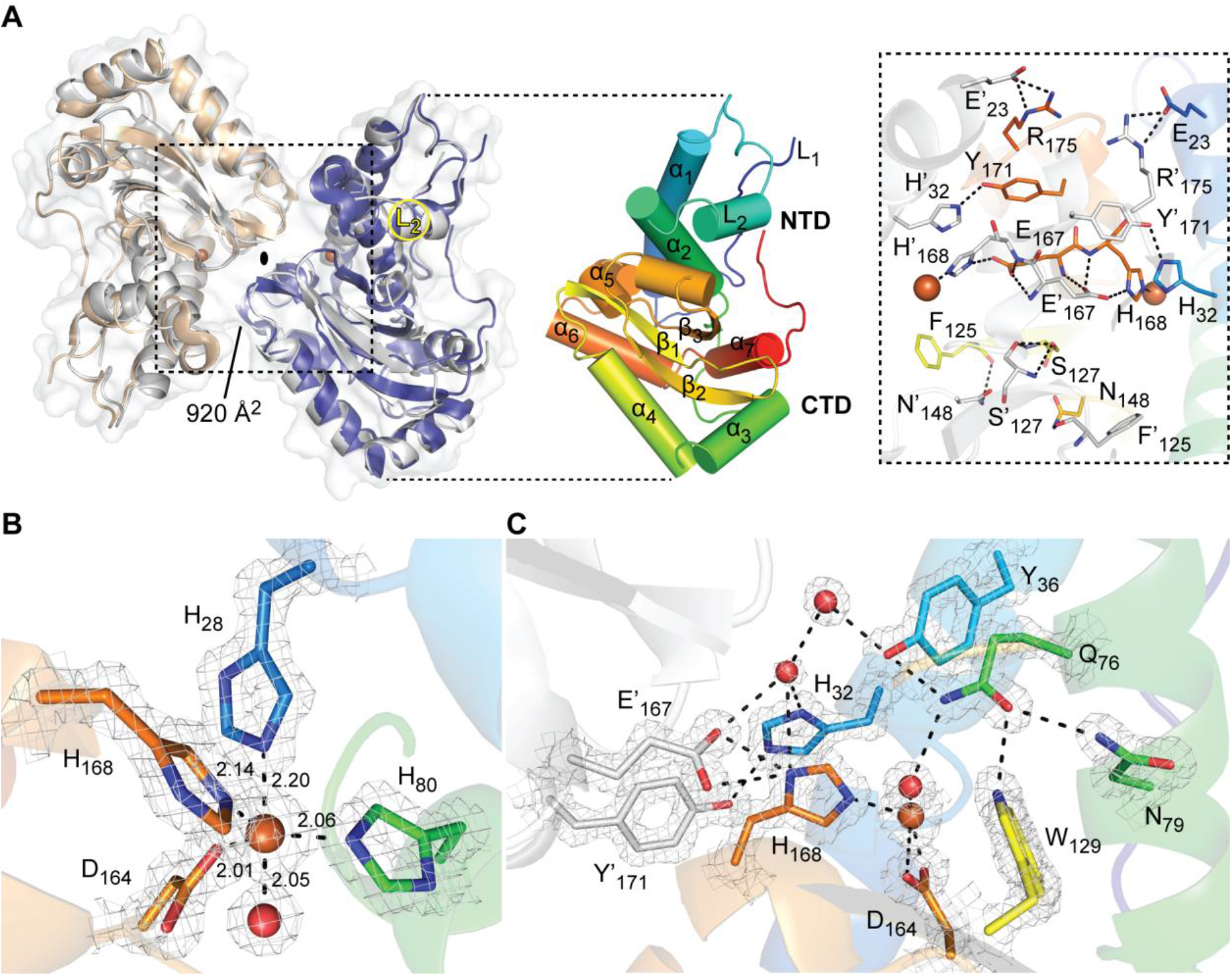
Crystal structure of ^AV3^SodB. (A) Although there is a single molecule of ^AV3^SodB per asymmetric unit, two contiguous molecules (in beige and blue, the interface area between them is informed in Å^2^) related by crystallographic symmetry (oval symbol) give rise to a dimer of the protein which presents a quaternary structure similar to that observed for the *E. coli* FeSOD (PDB code 1ISA, in Gray, superimposed to ^AV3^SodB) (on the left). Proteins are shown in ribbon representation. The surface of ^AV3^SodB is depicted in gray. On the right, a monomer of ^AV3^SodB is shown in rainbow colours, with helices represented as cylinders. Secondary structure elements conserved in FeSODs are indicated (see also Fig. 2A). NTD, N-terminal domain; CTD, C-terminal domain. An inset of the interface between ^AV3^SodB molecules in a protein dimer is shown. Residues involved in intermolecular hydrogen bonds or salt bridges are depicted in stick representation. Iron ions are shown in this and subsequent panels as orange spheres. (B) Active site of ^AV3^SodB. Metal ligands are represented as sticks. Interatomic distances are informed (in Å). One of the axial ligands is a water molecule (red sphere). (C) A conserved network of hydrogen bonds involving active site residues. Interatomic interactions are depicted as dashed lines. The mesh in panels (B) and (C) corresponds to the crystallographic *2mFo–DC* electron density map contoured to 2.0 σ.

**Table 1.** X-ray diffraction data collection and refinement statistics. One protein crystal was employed for structure determination. Values in parentheses are for the highest-resolution shell.

|  | **AV3SodB**<br>**(PDB 7SBH)** |
| --- | --- |
| **Data collection** |  |
| Space group | C222<sub>1</sub> |
| Cell dimensions |  |
| *a*, *b*, *c* (Å) | 70.93 87.02 75.61 |
| $\alpha$, $\beta$, $\gamma$ (°) | 90.00 90.00 90.00 |
| Resolution range (Å) | 28.54 - 1.34 (1.37 - 1.34) |
| *R*<sub>merge</sub> | 0.062 (0.749) |
| *R*<sub>pim</sub> | 0.026 (0.360) |
| *I*/σ*I* | 18.9 (2.4) |
| Completeness (%) | 99.7 (94.1) |
| Multiplicity | 6.6 (5.1) |
| **Refinement** |  |
| Resolution (Å) | 26.85 - 1.345 (1.393 - 1.345) |
| No. reflections | 51,972 (4,958) |
| *R*<sub>work</sub>/*R*<sub>free</sub> | 0.144/0.162 |
| Protein residues | 208 |
| Ligand molecules | 2 |
| No. atoms | 2,050 |
| Protein | 1,666 |
| Ligands | 32 |
| Water | 352 |
| Wilson B-factor (Å<sup>2</sup>) | 12.96 |
| Average B-factors (Å<sup>2</sup>) | 17.96 |
| Protein | 14.93 |
| Ligands | 42.79 |
| Water | 30.09 |
| R.m.s. deviations |  |
| Bond lengths (Å) | 0.012 |
| Bond angles (°) | 1.15 |
| Ramachandran |  |
| Favoured (%) | 97.57 |
| Allowed (%) | 2.43 |
| Outliers (%) | 0 |

The active site of ^AV3^SodB, located close to the dimer interface, is composed of conserved residues among FeSODs (Fig. 2A and Fig 5B). *mFo-DFc* electron density maps clearly revealed the presence of a metal ion bound at the active site, which, based on our bioinformatic analysis, was modelled as an iron ion. The metal ion is coordinated in a distorted trigonal bipyramidal geometry with the side chain of His28 and a solvent molecule as axial ligands and the side chains of His80, Asp164 and His168 as equatorial ligands (Fig. 5B). Residues His28 and His80 are located in helices α1 and α2, respectively, of the N-terminal domain of ^AV3^SodB, whereas Asp164 and His168 are located in the strand β3 of the C-terminal domain and the loop that follows it (Fig. 2A and Fig. 5B). ^AV3^SodB shares with *E. coli* FeSOD six out of seven residues at or near the active site that have been identified as specificity signature positions for metal ion use^10^.

The OH^-^/H_2_O ligand hydrogen bonds Asp164 and the active site Gln76 (Fig. 5C). This OH^-^/H_2_O, believed to be OH^-^ in oxidized FeSODs and H_2_O in the reduced enzymes^10^, is connected to bulk water *via* a conserved hydrogen bonding network that begins with Gln76 and continues with residues Tyr36 and His32. The side chain of Gln76 is stabilized by hydrogen bonding interactions with residues Tyr36, Asn79 and Trp129. Additional conserved hydrogen bonds link the active site to the interface between ^AV3^SodB monomers, including a hydrogen bond between His32 of one monomer and Tyr171 of the other, and a hydrogen bond between His168 of one monomer and Glu167 of the other.

Interestingly, ^AV3^SodB crystallized in the presence of flavin mononucleotide (FMN), added as an additive in homemade crystallization conditions. The isoalloxazine ring, clearly evident in *mFo-DFc* maps, is buried in a deep pocket formed by helix α2 in the N-terminal domain, loops L1 and L2 and the most C-terminal segment of the protein (Fig. S1). Notably, similar to the FeSOD from *H. pylori*^45^, the C-terminus of ^AV3^SodB is longer than for other related enzymes (Fig. 2A). The isoaloxazine ring of the FMN molecule is located at 12.4 A of the metal ion and it is possible to estimate an electron transfer path^46^ between them through the His33 and His80 residues (Fig. S1). Electronic absorption measurements after incubating ^AV3^SodB with FMN and removing the excess cofactor by size exclusion chromatography did not, however, give clear evidence of a binding event in solution (not shown). Furthermore, it is not evident what biological role the binding of FMN to this enzyme could have. To our knowledge, the binding of FMN to a superoxide dismutase has not been previously reported. Although the binding of FMN to ^AV3^SodB in the crystal is very intriguing, this could simply be an artifact and any further exploration of this observation will require specific experiments.

### ^AV3^SodB is a cytosolic enzyme whereas ^AV3^SodC is directed to the bacterial periplasm and loaded into OMVs

To determine the subcellular localization of ^AV3^SodB and ^AV3^SodC, we obtained polyclonal antibodies against the recombinant proteins and used them to perform Western Blot analyses of *Acinetobacter sp*. Ver3 cell fractions. Four different fractions were prepared: cytosol, soluble periplasmic fraction, insoluble periplasmic fraction and outer membrane vesicles (OMVs). ^AV3^SodB was detected in the bacterial cytosol whereas ^AV3^SodC was additionally found in the insoluble periplasmic fraction (Fig. 6A). Consistent with the subcellular localization of ^AV3^SodC, its N-terminal sequence comprises a ^(−3)^[LVI]^(−2)^[Xaa]^(−1)^[Yaa]^(+1)^C motif (Fig. 2C) that is a hallmark of bacterial lipoprotein attachment sites^41,42^, suggesting that ^AV3^SodC could be associated with the membrane.

**Figure 6.**
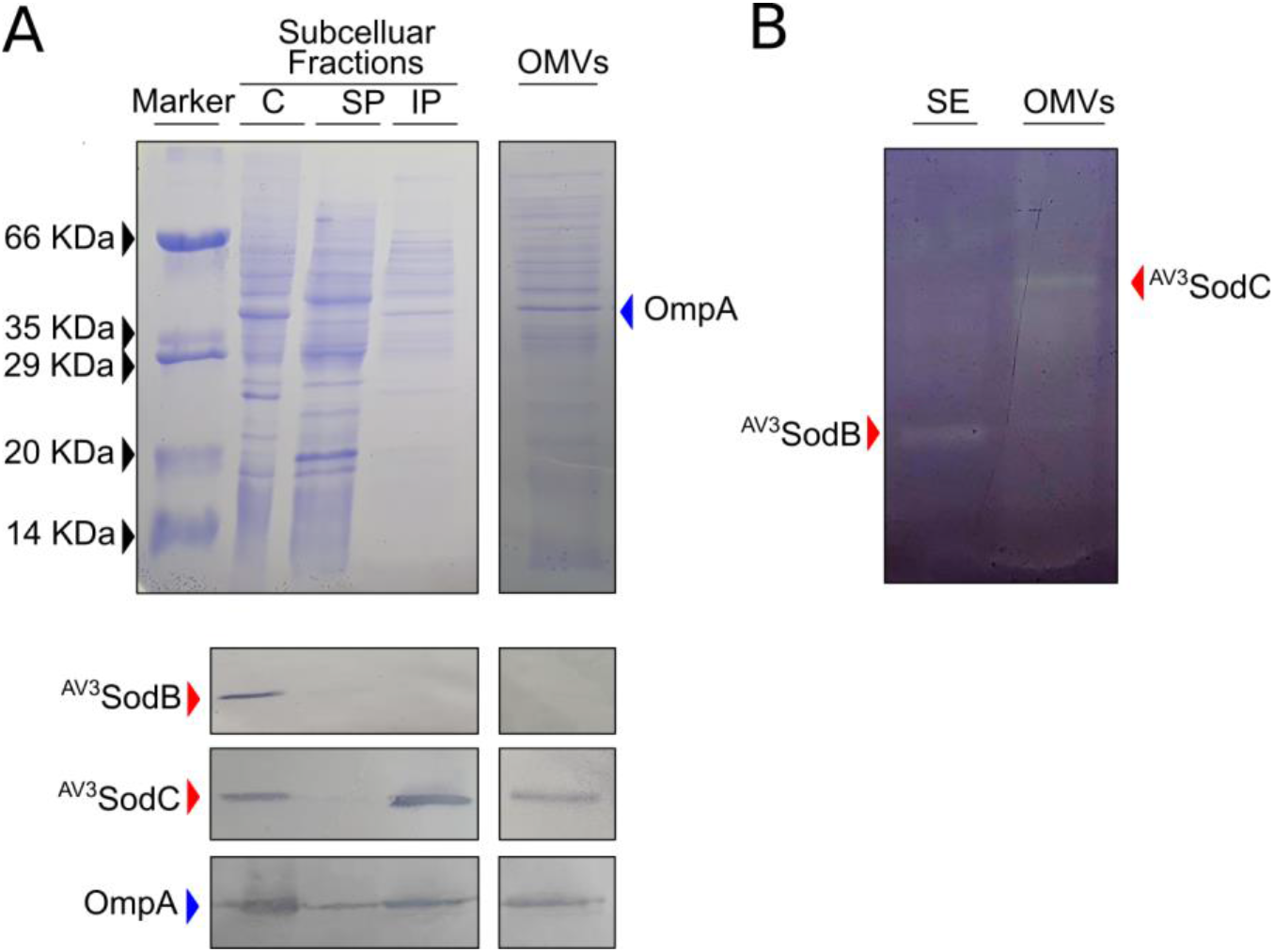
^AV3^SodB and ^AV3^SodC subcellular localization. (A) Coomassie stained SDS-PAGE (upper panel) and Western Blot assay (lower panel) of cytosolic (C), soluble and insoluble periplasmic (SP and IP, respectively) fractions (7 μg of total proteins), and OMVs (15 μL from a 500X culture concentrate) obtained from *Acinetobacter* sp. Ver3. Specific antibodies raised against ^AV3^SodB and ^AV3^SodC (red arrows) were used. The detection of OmpA (blue arrow) with specific antibodies against the *A. baumannii* protein was used as a control; OmpA has been shown to be loaded into OMVs^64^. (B) In-gel assessment of SOD activity present in a soluble extract (SE) (1.5 μg of total proteins) and OMVs (15 μL from a 500X culture concentrate) of *Acinetobacter* sp. Ver3.

Since the CuZnSOD from *A. baumannii* ATCC 17978 has been detected in OMVs^47^, we investigated whether this was also the case for ^AV3^SodC. Indeed, ^AV3^SodC was found in OMVs obtained from *Acinetobacter sp*. Ver3 cultures (Fig. 6A). Notably, ^AV3^SodC is active when located in OMVs (Fig. 6B).

### Differential expression of *sodB* and *sodC* in response to pro-oxidant challenges

In order to further explore the physiological roles of ^AV3^SodB and ^AV3^SodC, we studied the changes in *sodB* and *sodC* expression in *Acinetobacter* sp. Ver3 cells subjected to pro-oxidant challenges. qRT-PCR analyses were performed from total RNA samples obtained 10 and 30 min after the exposure of cells to MV, H_2_O_2_, UV radiation and blue light. The latter treatment was carried out because blue light acts as a pro-oxidant agent in bacteria by generating ROSs^48^. On the other hand, blue light induces catalase activity in *Acinetobacter baumannii*^49^, and thus could play an important role in the defence against oxidative stress.

The expression of *sodC* increased about 2-fold after a 30 min blue light challenge, but no significant changes were observed under the rest of the pro-oxidant conditions (Fig. 7A). In contrast, *sodB* transcript levels remained unaltered under the conditions tested. To determine whether these results showed a correlation at the protein level, exponentially growing cell cultures were exposed to blue light and protein extracts were then obtained. Soluble fractions as well as OMVs were analysed by SDS-PAGE followed by anti-^AV3^SodC immunostaining (Fig. 7B). A net accumulation of ^AV3^SodC was observed in OMVs after the blue light treatment.

**Figure 7.**
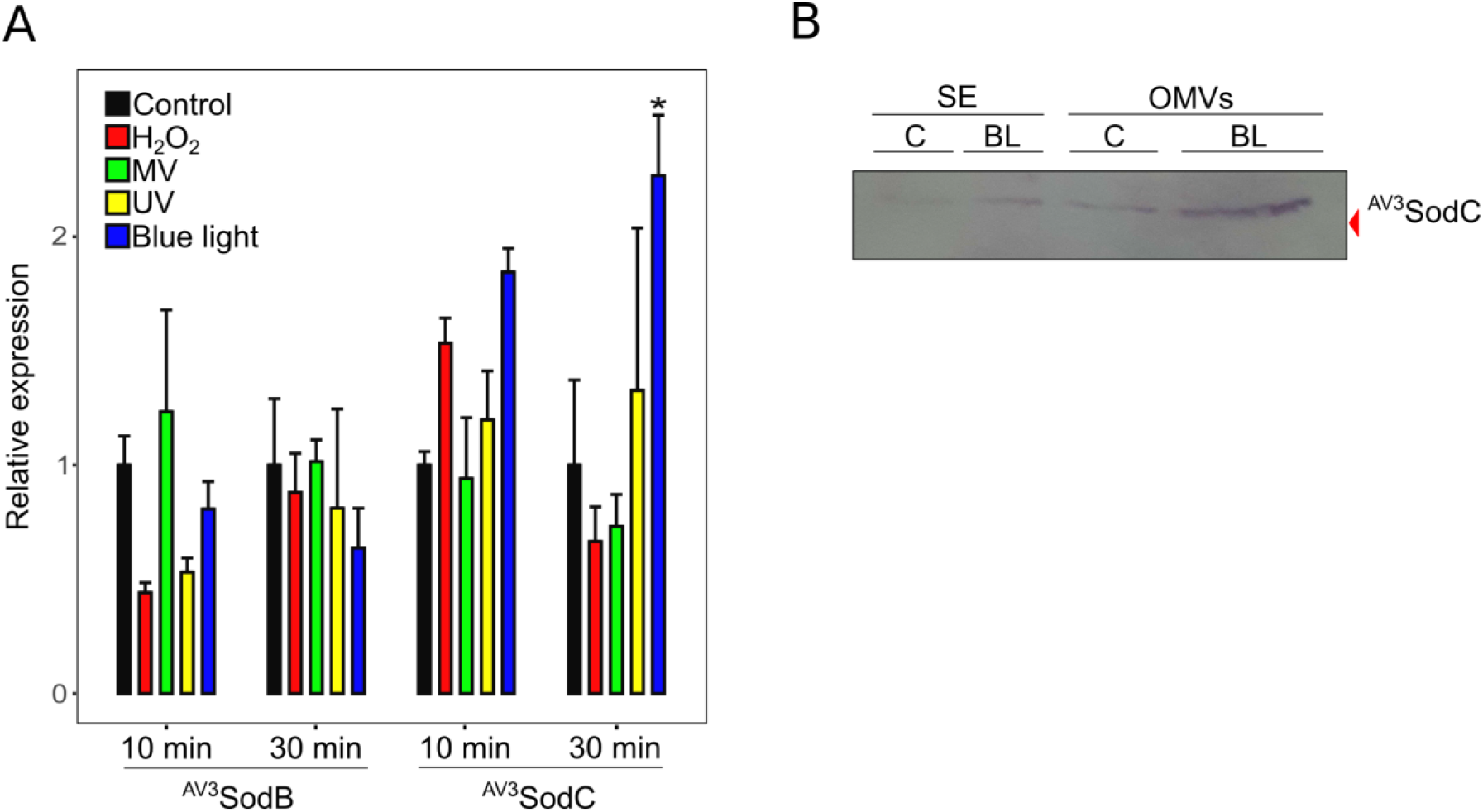
*sodB* and *sodC* response to pro-oxidant challenges. (A) Relative levels of *sodB* and *sodC* transcription in untreated *Acinetobacter sp*. Ver3 cells (black bars) or after exposure to 1 mM H_2_O_2_ (red bars), 0.5 mM Methyl Viologen (MV) (green bars), 900 J.m^-2^ ultraviolet (UV) (yellow bars) or blue light (blue bars). The mean for the housekeeping genes *recA* and *rpoB* was used as a normalizer. Asterisks indicate significant differences among control and treated samples, as determined by analysis of variance (ANOVA) and Tukey’s multiple comparison test. In each case, the average value and the standard deviation of four biological replicates are shown. (B) Anti-^AV3^SodC immunoblot of a soluble extract (SE, 7 μg of total proteins) and OMVs (15 μL from a 500X culture concentrate) of *Acinetobacter* sp. Ver3 grown over 26 h in the presence of blue light (BL, 5 μmol.m^-2^.s^-1^). A control (C) culture was also included.

### Prevalence of SOD-encoding genes in the *Acinetobacter* genus

In order to explore the diversity of SOD-encoding genes in the *Acinetobacter* genus, we assembled a local database with 314 publicly available complete genomes, comprising 53 *Acinetobacter* defined species as well as 21 strains corresponding to unassigned species (Table 2). BLASTp searches were then performed using the sequences of the enzymes FeSOD (ABO12765.2)^19^ and CuZnSOD (ABO13540.2) from *A. baumannii* ATCC 17978 as queries. Following the strategy described in the Methods section, we found 677 proteins (Table S3), and by inspecting the SOD-encoding genes in the selected *Acinetobacter* strains, we distinguished three genotypes (Table 2):

i. Genotype 1 (1 FeSOD + 1 CuZnSOD; 83.4 % of the strains). This group contains representatives of 28 species, but it is remarkably enriched in those that have been associated with nosocomial infections, such as *A. baumannii, A. pittii* and *A. nosocomialis*. It also includes 17 unassigned strains, such as *Acinetobacter sp*. Ver3. CuZnSOD enzymes in this group contain a predicted lipoprotein signal peptide and could thus be translocated to the periplasm or the extracellular space similarly to ^AV3^SodC. We also include in this group three *A. baumannii* strains (921, 3207, 7835), bearing two FeSOD encoding genes, which were denominated *sodB1* and *sodB2*.
ii. Genotype 2 (1 FeSOD + 1 MnSOD + 1 CuZnSOD; 10.2 % of the strains). This class encompasses a total of 18 *Acinetobacter* species and 3 unassigned strains. MnSODs in 27 out of 32 strains of this group are predicted to be cytosolic enzymes, with the exception of one *A. beijerinckii* and three *A. junii* strains, which encode MnSOD variants with a predicted signal peptide without a lipidation motif (Fig. S2). Interestingly, *A. bereziniae* XH901 encodes MnSODs of the two types, comprising a total of 4 SOD encoding-genes in the genome. Similar to genotype 1, CuZnSOD enzymes in this group contain a predicted lipoprotein signal peptide.
iii. Genotype 3 (1 FeSOD + 1 or 2 MnSOD; 6.1 % of the strains). This group contains 7 species comprising a total of 18 strains, with *A. haemolyticus* the most prevalent species (11 strains). One strain corresponding to an *Acinetobacter* unassigned species is also included in this group. Remarkably, all strains encode a MnSOD enzyme containing a putative signal peptide without a lipidation motif (Fig. S2), which could be directed to the periplasmic space to play a CuZnSOD-like role. Twelve strains additionally bear a putative cytosolic MnSOD.

**Table 2.**
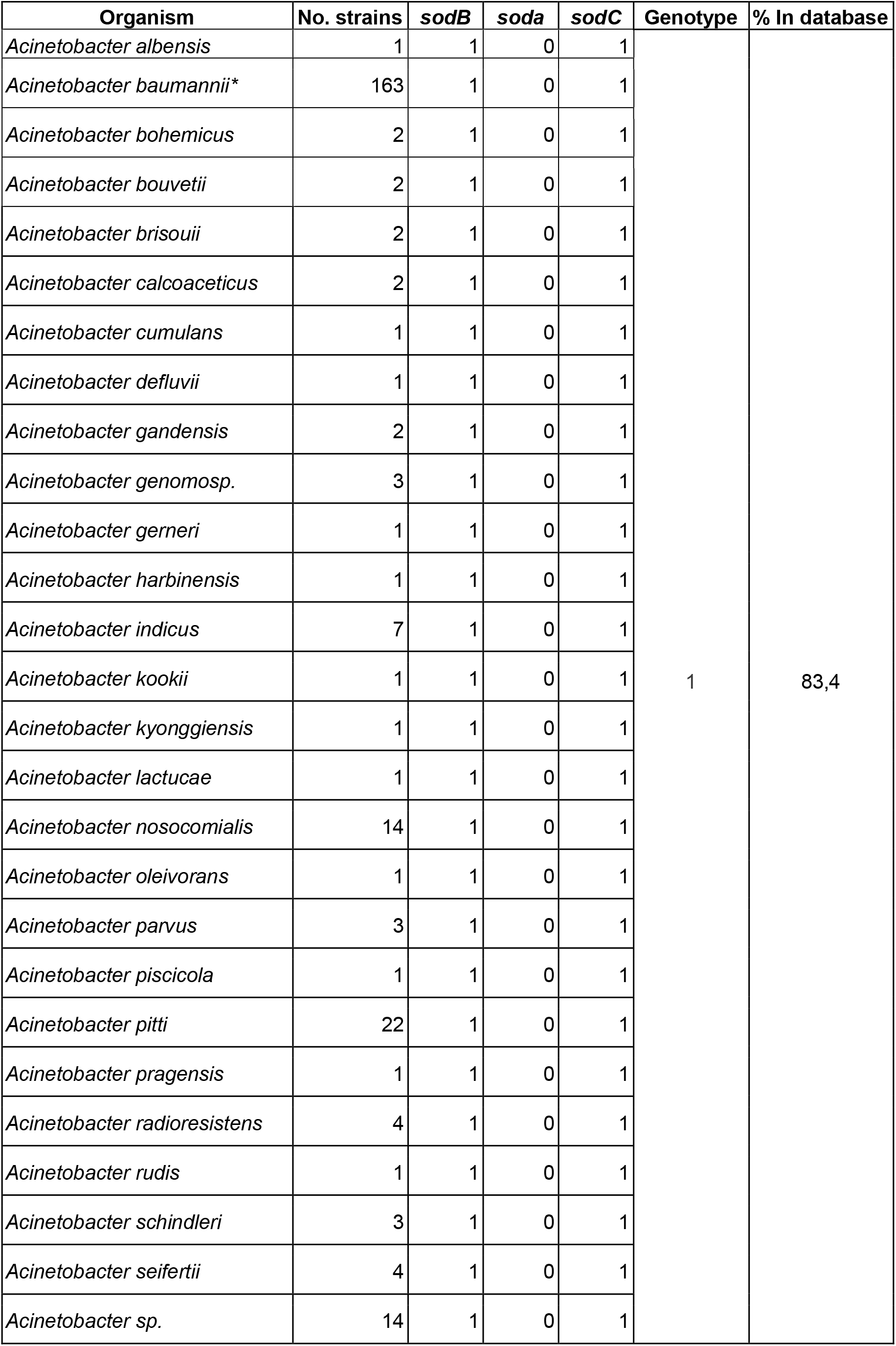
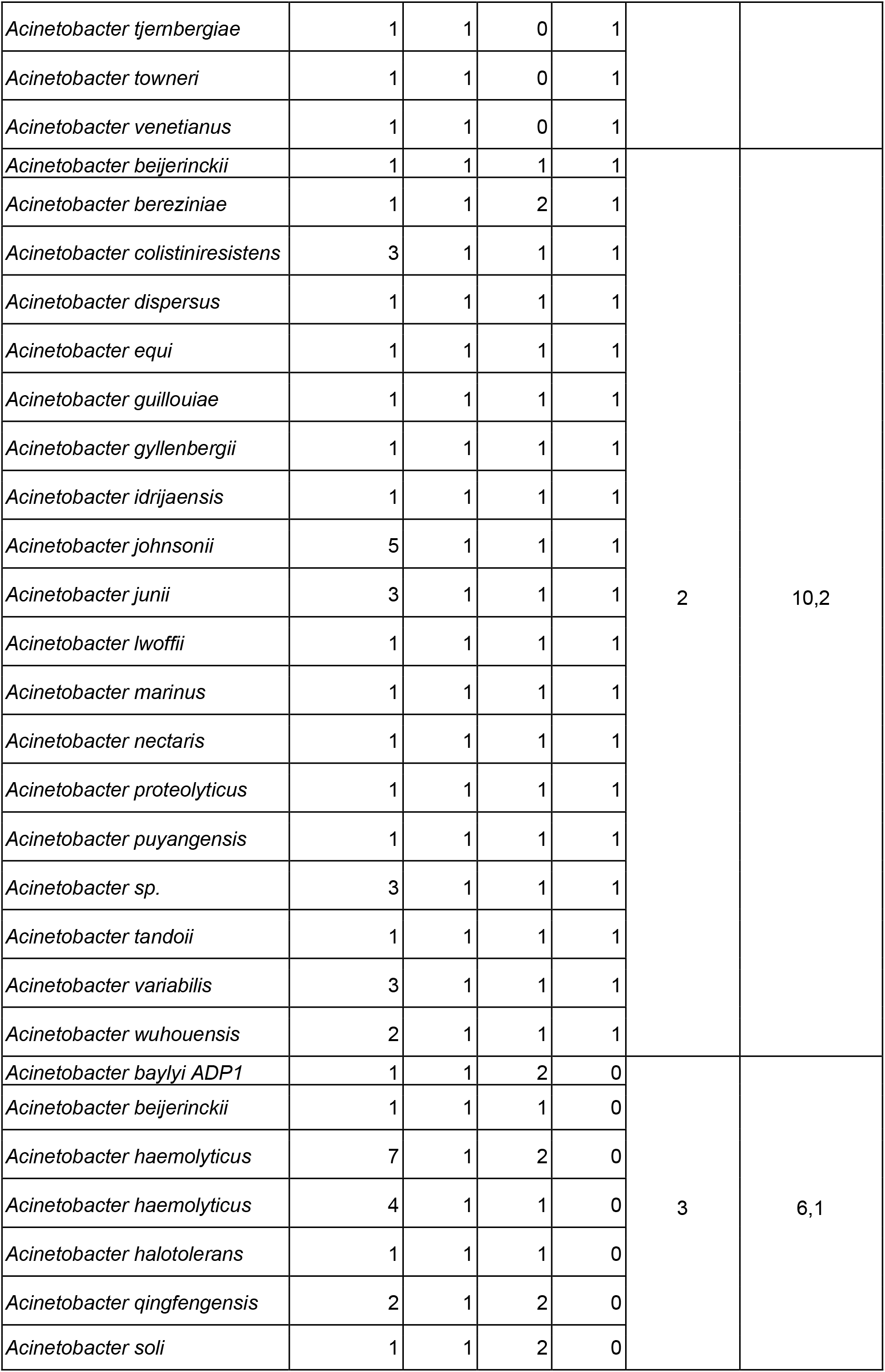
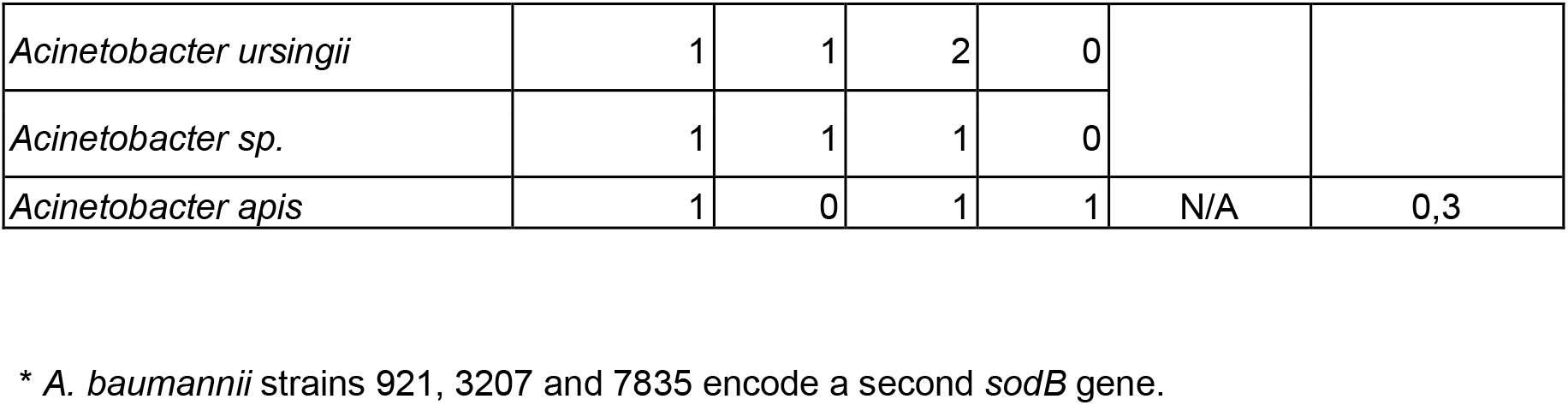
Prevalence of *sod* genes in the genus *Acinetobacter*.

An uncategorized case is that of the species *A. apis*, which encodes 1 CuZnSOD and 1 MnSOD (Table 2), but lacks a FeSOD-encoding gene. The MnSOD is predicted to be cytosolic (Fig. S2).

## DISCUSSION

SOD enzymes are ubiquitous in biology, so understanding the principles that govern their kinetic parameters, the use of metal cofactors and the subcellular localization has diverse implications. From a biotechnological point of view, the characterization of SODs isolated from extremophiles is attractive since these enzymes usually work under advantageous conditions in industrial processes. From a medical perspective, SODs are part of the bacterial defence against oxidative stress and by detoxifying the superoxide produced by the mammalian immune system in response to infection they play a crucial role in pathogenesis. In this work we carried out a biochemical and structural characterization of ^AV3^SodB and ^AV3^SodC, the SOD enzymes encoded by the extremophile *Acinetobacter sp*. Ver3. Such analyses were accompanied by an investigation of gene expression and the subcellular localization of the proteins. Finally, we assessed the prevalence of different SOD types in the genus *Acinetobacter*, in which not only free-living species are found, but also important pathogens.

The specific activity of ^AV3^SodB (6,600 ± 200 U/mg) resulted similar to that of *E. coli* FeSOD (6,700 U/mg)^50^, although ^AV3^SodB (Fig. 4A and C) showed a higher thermal stability and activity in a broader pH range, comparable to the reported properties of the iron form of the cambialistic Fe/MnSOD from *Propionibacterium shermanii*^51^. On the other hand, ^AV3^SodC^-p^ (^AV3^SodC devoid of its N-terminal sequence) presented a lower specific activity (1,800 ± 200 U/mg) than *E. coli* CuZnSOD (3,700 U/mg)^52^ and ^AV3^SodB. Besides, ^AV3^SodC^-p^ activity was more susceptible to heat inactivation and extreme pHs than ^AV3^SodB (Fig. 4A, B and C). The activity of ^AV3^SodB and ^AV3^SodC^-p^ was also tested in the presence of different potential inhibitors (Fig. 4D). BME in a concentration of up to 10 mM did not alter the activity of either of the two enzymes. In the case of ^AV3^SodB, this is consistent with the absence of Cys residues in the protein sequence; on the other hand, although ^AV3^SodC^-p^ has three Cys residues, these results do not support a modulation of the enzyme activity in response to changes in their oxidation state. Of all the other chemical agents tested, including a chelating agent, two detergents, two denaturing agents and an organic solvent, only the addition of 50% ethanol to ^AV3^SodB resulted in an almost complete loss of activity; on the other hand, for all other treatments the two enzymes retained at least 50% of the basal activity, maintaining more than 80-90% of it in most cases. These properties make *Acinetobacter sp*. Ver3 SOD enzymes attractive for use in industrial applications. Besides, we report the crystal structure of ^AV3^SodB to 1.34 Å resolution, one of the highest resolutions achieved for an enzyme of this type. ^AV3^SodB bears *ca*. 90 % amino acid identity with the equivalent enzyme in *A. baumannii*, an important target for drug design^19^. Thus, we present detailed biochemical and structural data relevant in oxidative stress processes in environmental as well as pathogenic species.

We provide evidence that ^AV3^SodB is a cytosolic enzyme whereas ^AV3^SodC is also found in the periplasmic space (Fig. 6A). What is more, we show that ^AV3^SodC is loaded in an active state in OMVs (Fig. 6A and B), consistent with previous reports for *A. baumannii*^47^. The presence of ^AV3^SodC in OMVs suggests that this enzyme, in addition to playing a protective role in the periplasmic space, could also act by regulating the redox state in the extracellular microenvironment. Interestingly, while *sodB* transcriptional levels remained unaltered, a blue light treatment induced a twofold increase in the expression of *sodC* (Fig. 7A), which was accompanied by a higher amount of the protein detected in OMVs (Fig. 7B). Previous studies concluded that blue light triggers the production of ROSs and leads to cell damage^53^, being able to modulate the metabolism, virulence, motility and even the tolerance to antibiotics of *Acinetobacter* species^49,54,55^. Altogether, these results indicate that SodC activity is critical for these bacteria to mount an effective defence against oxidative damage.

According to our bioinformatic analysis, 52 out of 53 *Acinetobacter* species analysed contain at least one *sodB* gene encoding a cytoplasmic FeSOD (Fig. 8A, B and C). The only exception was *A. apis* HYN18(T), a strain isolated from the intestinal tract of a honeybee, which shows a relatively small 2.41 Mbp genome^56^. We hypothesize that the adaptation of this strain to the host intestinal tract has led to a genomic reduction, with the consequent loss of the *sodB* gene. Still, *A. apis* conserves a *sodA* gene coding for a cytoplasmic MnSOD, highlighting the role of cytoplasmic SODs in response to oxidative stress. On the other hand, 46 out of 52 *Acinetobacter* species encode a periplasmic CuZnSOD (Genotypes 1 and 2, Fig. 8A and B), while seven species (Genotype 3, Fig. 8C) encode instead a MnSOD with a putative signal peptide (Fig. S2). Such MnSODs, lacking a lipidation motif, could perform functions similar to those of CuZnSODs in the bacterial periplasm, although they would not be directed to (and thus would not act on) OMVs. The presence of a periplasmic SOD in all *Acinetobacter* strains under study suggests that such a variant is necessary for crucial protective roles in the periplasmic space. This hypothesis is supported by several works that have demonstrated the participation of periplasmic SODs in the resistance to the respiratory burst elicited by the human immune system in response to pathogens^57^.

**Figure 8.**
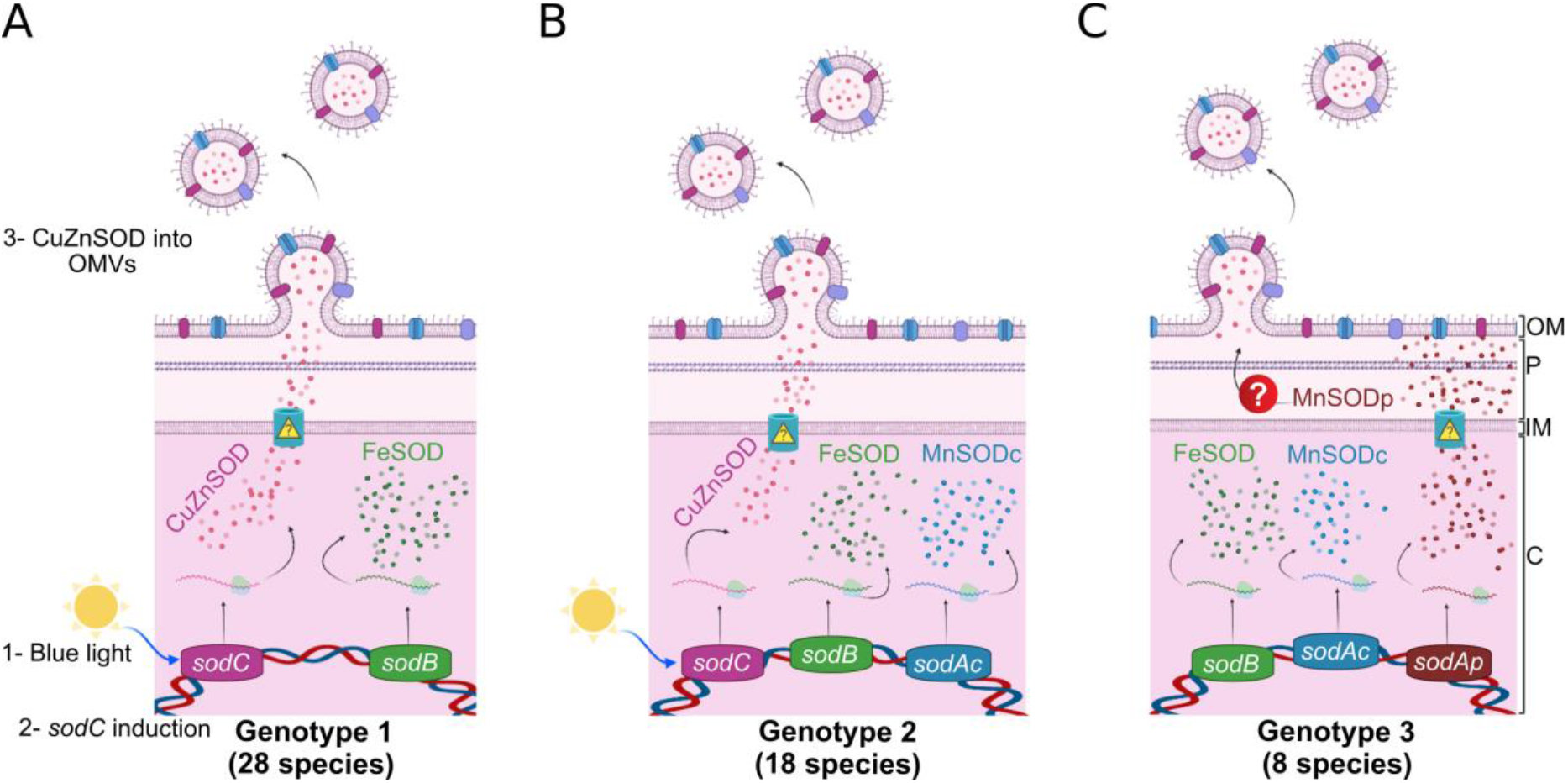
Proposed model schematizing the subcellular localization of SODs encoded by *Acinetobacter spp*. according to their genotype. (A) Strains with a type 1 genotype harbour *sodB* and *sodC* genes, coding for a FeSOD and a CuZnSOD, which are located in the cytosolic (C) and periplasmic (P) spaces, respectively. The CuZnSOD could also be loaded into OMVs, as shown for *Acinetobacter* sp. Ver3 strain. (B) Strains with a type 2 genotype additionally harbour a *sodAc* gene, coding for a cytosolic MnSOD (cMnSOD). (C) Strains with a type 3 genotype bear *sodB* and *sodAp* genes, coding for a FeSOD and a periplasmic MnSOD (pMnSOD), respectively. In 12 out of the 19 strains in this group, a *sodAc* gene is also present. *One strain of *A. beijerinckii* is present in each of these groups. IM: inner membrane; OM: outer membrane.

Most pathogenic *Acinetobacter* species, included within the ABC complex, code for only two SODs: a cytosolic FeSOD and a periplasmic CuZnSOD (Genotype 1, Fig. 8A). On the other hand, environmental strains, which are closer to the root of the *Acinetobacter* phylogenetic tree^58^, have more diverse genotypes regarding encoded SODs (Genotypes 1 to 3, Fig. 8A, B and C). According to previous findings for SODs in the genus *Staphylococcus*, it has been proposed that ancestral species often harbour MnSODs while the more recent ones, usually linked to infectious processes, have evolved cambialistic Fe/MnSODs due to the low bioavailability of manganese ions in the host^39^. The differences observed among genotypes in *Acinetobacter* can in fact be explained similarly. Thus, nosocomial *Acinetobacter* species might have lost cytoplasmic MnSODs in favour of FeSODs encoding genes; even though the bioavailability of iron ions is also scarce, *Acinetobacter* species harbour an arsenal of mechanisms for their scavenging^59,60^. On the other hand, periplasmic MnSODs in *Acinetobacter* might have been replaced by CuZnSODs during the evolution of pathogenic species.

The identification of *sodB2*, a gene possibly acquired by horizontal gene transfer (Fig. S3) and which encodes an additional FeSOD in three clinical strains of *A. baumannii* (Table 2), supports the notion that this SOD type is prevalent in modern strains. The *sodB2* genes are located next to a *GIsul2* genomic island^61,62^, which contains a *sul2* gene that confers resistance to sulphonamides as well as genes that confer resistance to tetracycline and chloramphenicol. Given that exposure to antibiotics generates ROSs and thus produces oxidative stress in *Acinetobacter*^63^, it is tempting to speculate that SOD activity acts in conjunction with antibiotic resistance mechanisms to counteract the deleterious effects of antimicrobial agents. In favour of this hypothesis, a *sodB1* mutation in *A. baumannii* ATCC17978 leads to increased susceptibility to antibiotics^19^.

Our bioinformatic analysis support the critical role played by SOD enzymes in the *Acinetobacter* genus, which thereby can be proposed as new therapeutic targets in pathogenic species such as *A. baumannii*. The high-resolution structural data presented here provides instrumental information for the design of SodB-specific inhibitors to control nosocomial infections caused by opportunistic bacteria. Additionally, the biochemical information reported for the SOD enzymes studied may be relevant for future applications in food, agriculture or industrial processes. In summary, this multidisciplinary approach serves to frame future studies on the role of SOD enzymes in the defence of pathogenic or environmental Acinetobacter species against oxidative stress.

## Supporting information

PDB Data

## DATA AVAILABILITY

Structure factors and atomic coordinates were deposited in the Protein Data Bank under the accession code 7SBH.

## ACKNOWLEDGEMENTS

This paper is dedicated to the memory of Dr. Cortez, a distinguished colleague and beloved friend.

## FUNDING

This work was supported by grants from the Agencia Nacional de Promoción de la Investigación, el Desarrollo Tecnológico y la Innovación (Agencia I+D+i, Argentina) received by N.C. (PICT 2015-1492); B.A.S. is a doctoral fellow of the Consejo Nacional de Investigaciones Científicas y Técnicas, Argentina (CONICET); M.G.S. is a former doctoral fellow of CONICET; D.A. and M.N.L. are researchers of CONICET; G.D.R. is a former researcher of CONICET.

## AUTHOR CONTRIBUTIONS STATEMENT

B.A.S. designed and performed experiments and participated in the interpretation of results. M.G.S. performed RT-qPCR assays. D.A. designed and performed crystallization trials. M.N.L. designed the crystallographic experiment, collected X-ray diffraction data, solved the crystal structure of ^AV3^SodB, interpreted results and wrote the paper. G.D.R. performed bioinformatics analyses and wrote the paper. N.C., M.N.L. and G.D.R. developed concepts and designed the work. All authors read and approved the final manuscript.

## CONFLICT OF INTERESTS STATEMENT

The authors declare no conflict of interest.

## SUPPLEMENTARY TABLES

**Table S1. Strains and plasmids used in this work.**

**Table S2. Oligonucleotides used in this work.**

**Table S3. Search of SODs encoded by *Acinetobacter spp*.**

## SUPPLEMENTARY FIGURES

**Figure S1. A FMN binding site in ^AV3^SodB.**

**Figure S2. Sequence alignment of MnSODs in *Acinetobacter spp.*.**

**Figure S3. Genomic localization of the *sodB2* gene.**

## Notes

### Competing Interest Statement

The authors have declared no competing interest.

