## Supplementary Data for "Coping with oxidative stress in extreme environments: the distinctive roles played by *Acinetobacter* sp. Ver3 superoxide dismutases"

**SUPPLEMENTARY MATERIAL**

**Table S1. Strains and plasmids used in this work.**

| | **Description** | **Reference** |
|---|---|---|
| Strains | | |
| *Acinetobacter sp. Ver3* | | Ordoñez *et al.,* 2009 |
| *E. coli* | | |
| DH5a | Φ80 *lacZ*DM15 *rea*A1 *end*A1 *gyr*A96 *thi*-1 *hsd*R17 *sup*E44 *rel*A1 *deo*RD(*lac*ZYA-*arg*F) U169 F | Woodcock *et al.,* 1989 |
| QC774(DE3) | $F^-$ *ompT* ($r_B^-$ $m_B^-$) *gal dcm* λ(DE3) pLysS ($Cm^r$ ) | Carlioz & Touati 1986 |
| Plasmids | | |
| pET32m | Expression vector, $Amp^R$, derived from pET32b, with a deletion on a enterokinase restriction site | Tabares *et al.,* 2006 |
| pET3228 | *oriR*(ColE1) *oriR*(f1) $Amp^R$ | Tabares *et al.,* 2006 |
| pESodB | pET3228 derivative expressing $^{AV3}$SodB | This work |
| $pESodC^{-p}$ | pET32m derivative expressing $^{AV3}$SodC without the sequence encoding the signal peptide | This work |
| pGEM®T-Easy | Cloning vector | Promega® |

**Table S2. Oligonucleotides used in this work.**

| **Primers** | **Sequence (5'–3')** |
|---|---|
| FMSOD3228F | 5'-CCATCCATGGCAACGATTACTTTACCAGCTCTTCC-3'[a] |
| FMSOD3228R | 5'-ACGGGAGCTCAATCTTATTTTTCTACGCCAGCTTCTTG-3'[b] |
| CSOD$^{sp}$32F | 5'-AGAAGGATCCGCAACGCAAAATACATCTGCATC-3'[c] |
| CSOD$^{sp}$32R | 5'-CCATCTCGAGGCTTAGCGTATAACACCACATGC-3'[d] |
| qFMSODF | 5'-AGGTATCTTCAACAACGCAGC-3' |
| qFMSODR | 5'-AGCAACTAACCAAGCCCAAC-3' |
| qCZSODF | 5'-ACCTGGATATCACGGGTTCC-3' |
| qCZSODR | 5'-CGCGTCAACATTTAACACTGG-3' |
| qrecAF | 5'-CTCAATATGCTCGCAAACTTGG-3' |
| qrecAR | 5'-GGTTAAGGCTGCTACAGAATCG-3' |
| qrpoBF | 5'-TGCAAACACGGTTCTTAGCC-3' |
| qrpoBR | 5'-CACCTGGACGCATTACCTTG-3' |

Underlined sequences correspond to recognition sites for restriction enzymes: [a]*Nco*I, [b]*Sac*I, [c]*Bam*HI, [d]*Xho*I, [e]*EcoR*I and [f]*Hind*III.

**Table S3. Search of SODs encoded by *Acinetobacter spp*.** Local Blast analysis of SOD proteins in *Acinetobacter spp.*, using the FeSOD and CuZnSOD from *A. baumannii* ATCC17978 as query.

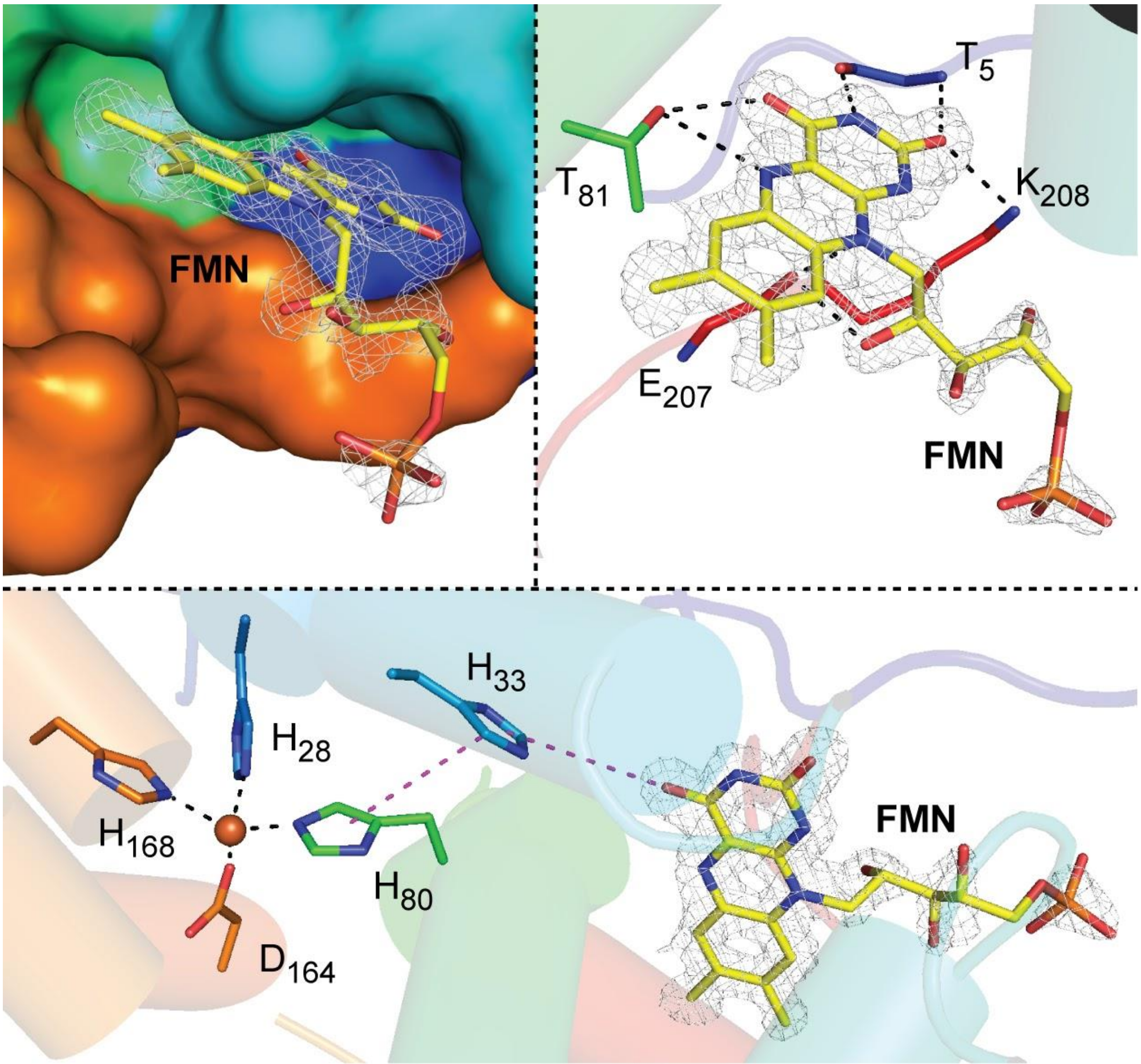


**Figure S1. A FMN binding site in $^{AV3}$SodB.** Upper panels: the binding pocket is shown in the protein in rainbow colours, in surface and ribbon representation. In the latter, the residues that establish polar interactions (black dashed lines) with the FMN are shown as sticks. The FMN molecule is also presented as sticks and the corresponding *2mFo–DFc* electron density (contoured to 1.0 σ) is displayed as a mesh. Bottom panel: relative position of the FMN molecule with respect to the active site of the enzyme. It is possible to estimate an electron transfer path (magenta dashed lines) (https://emap.bu.edu) between the FMN and residue His80 at the active site, which involves the His33. The iron ion is depicted as an orange sphere.

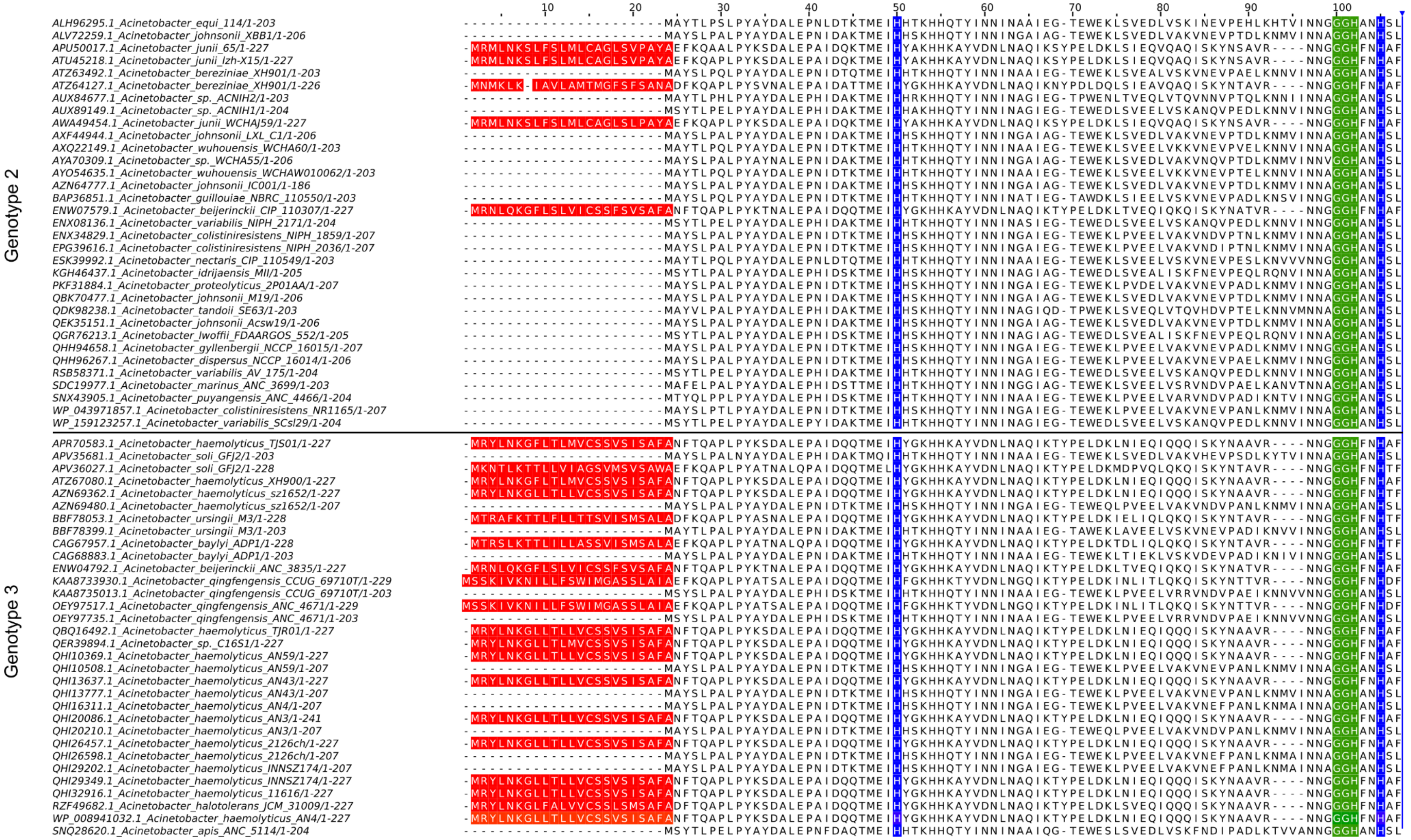


**Figure S2. Sequence alignment of MnSODs in *Acinetobacter spp.*.** In red, the putative signal peptides; in green, the characteristic MnSOD motif GGH; in blue, two first histidine residues from the metal binding site. Genotypes assigned in Table 2 are shown on the left. The C-terminal part of the proteins was intentionally deleted for a better visualization of the alignment.

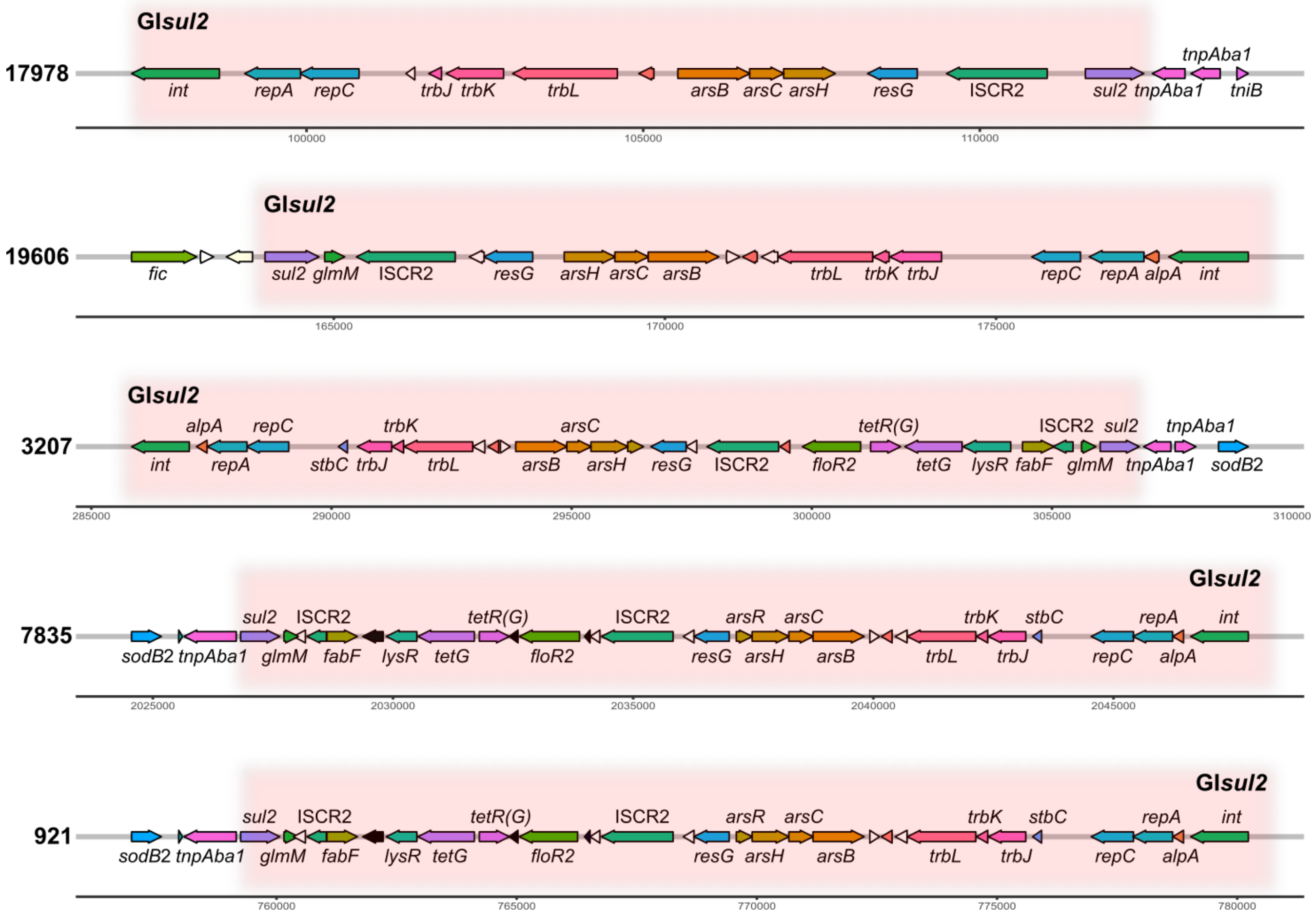


**Figure S3. Genomic localization of *sodB*2.** The *A. baumannii* strains 3207, 7835 and 921 encode a paralogous *sodB* gene (blue arrow), located close to a genomic island showing similarities to the GI*sul2* genomic island (pink box) found in *A. baumannii* ATCC17978 and ATCC19606 strains.
